# Mfn2 promotes NCLX-mediated Ca^2+^ release from mitochondria

**DOI:** 10.1101/2024.08.05.606704

**Authors:** Panagiota Kolitsida, Akash Saha, Andrew Caliri, Essam Assali, Alejandro Martorell Riera, Samuel Itskanov, Sean Atamdede, Catalina S. Magana, Björn Stork, Orian Shirihai, Israel Sekler, Uri Manor, Carla M. Koehler, Alexander M. van der Bliek

## Abstract

Mfn2 is a mitochondrial outer-membrane fusion protein that also functions as a tether between mitochondria and the ER. Here, we identify a previously unrecognized role for Mfn2 in promoting mitochondrial Ca^2+^ release via the Na^+^/Ca^2+^ exchanger NCLX. This function was uncovered through studies with the fungal toxin phomoxanthone A (PXA), which induces NCLX-dependent Ca^2+^ release by directly targeting Mfn2. Mfn2-dependent Ca^2+^ release through NCLX is similarly triggered by ROS in respiring cells treated with oligomycin or mitoPQ. ROS enhances Ca^2+^ release by strengthening the interaction between Mfn2 and NCLX, an interaction that also requires the mitochondrial outer-membrane protein SLC25A46. Together, these proteins coordinate mitochondrial fission and Ca^2+^ release to initiate mitophagy. The antioxidant N-acetylcysteine blocks ROS-induced mitochondrial fission, but inhibition of Ca^2+^ release with the NCLX inhibitor CGP37157 does not, indicating that ROS-driven fission is independent of Ca^2+^ release. In contrast, Ca^2+^ release is required for efficient mitophagy, as NCLX inhibition arrests this process at a later stage. We further show that Ca^2+^ promotes mitophagy through NEDD4-1, which is a Ca^2+^-responsive E3 ubiquitin ligase. Together, these findings connect mitochondrial ROS production to Ca^2+^ signaling, mitochondrial remodeling, and mitophagy, providing new insight into how mitochondrial dysfunction may contribute to neurodegenerative and metabolic disease.

## Introduction

Fusion between mitochondrial outer membranes is mediated by the mitofusins Mfn1 and Mfn2 in mammals (Labbe *et al*, 2014). These proteins are dynamin-related GTPases with transmembrane segments anchored in the mitochondrial outer membrane (Santel & Fuller, 2001). Mutations in either mitofusin impair mitochondrial fusion and they are partially redundant with the ability to form heteromeric complexes (Chen *et al*, 2003), but they also have distinct functions. Mfn1 supports stress-induced mitochondrial hyperfusion (SIMH) (Tondera *et al*, 2009), while Mfn2 localizes to sites of mitochondrial fission at mitochondria-ER contact sites (MAMs) (Karbowski *et al*, 2002) and functions as a tether that promotes mitochondrial-ER association (de Brito & Scorrano, 2008). Through this role as a tether, Mfn2 influences several MAM-associated processes, including transfer of coenzyme Q (Mourier *et al*, 2015) and phosphatidylserine (Hernandez-Alvarez *et al*, 2019), as well as broader metabolic functions (Sebastian *et al*, 2012).

Mfn2-dependent ER-mitochondria contacts are also important for Ca^2+^ homeostasis (Wang *et al*, 2025b; Yang *et al*, 2023). Ca^2+^ released from the ER is taken up by the mitochondrial Ca^2+^ uniporter (MCU) in the inner membrane (De Stefani *et al*, 2015), while Ca^2+^ efflux occurs primarily through the Na^+^/Ca^2+^ exchanger NCLX (Palty *et al*, 2010), which is regulated by PKA phosphorylation and membrane potential (Kostic *et al*, 2018; Kostic *et al*, 2015; Kostic & Sekler, 2019). Mitochondrial Ca^2+^ release can promote mitochondrial fragmentation via Ca^2+^/calmodulin-dependent protein kinase I and II mediated phosphorylation (Han *et al*, 2008; Xu *et al*, 2016), and calcineurin mediated dephosphorylation of Drp1 (Cereghetti *et al*, 2008). AMP-activated protein kinase (AMPK), a central energy sensor, further links metabolism to mitochondrial dynamics. Under low-ATP conditions, AMPK promotes fission by phosphorylating the mitochondrial Drp1 receptor MFF (Toyama *et al*, 2016). AMPK also phosphorylates Mfn2 at serine 442 (Hu *et al*, 2021), a modification important for mitochondrial remodeling during metabolic stress. Mutations in MFN2 are the most common cause of axonal peripheral neuropathy, Charcot-Marie-Tooth type 2 (CMT2) (Verhoeven *et al*, 2006; Zuchner *et al*, 2004), but it remains unclear which Mfn2 functions are most relevant to the disease.

While studying previously described effects of the fungal metabolite phomoxanthone A (PXA) on mitochondrial Ca^2+^ release (Bohler *et al*, 2018), we discovered that these effects are suppressed by Mfn2 gene mutation. We then show that Mfn2 cooperates with NCLX to promote Ca^2+^ release, a process hinted at by previous findings showing that Mfn2 can drive Ca^2+^ influx through NCLX instead of efflux in isolated mitochondria (Samanta *et al*, 2018). The outer-membrane protein SLC25A46 acts as an adaptor linking Mfn2 to NCLX; its deletion disrupts both mitochondrial Ca²⁺ release and Mfn2-NCLX association. ROS also induce Mfn2-dependent Ca^2+^ release via NCLX, prompting investigation of upstream regulatory steps and downstream physiological relevance of this pathway. We find that Oma1-mediated cleavage of Opa1 and PKA-mediated phosphorylation of NCLX both regulate ROS-induced Ca^2+^ release via NCLX. Downstream effects of ROS include AMPK-dependent fission and cytosolic Ca^2+^-dependent activation of E3 ubiquitin ligases, which are essential for mitophagy. Overall, these findings reveal a mechanism relevant to neurodegenerative disease: mutations in these proteins cause defective mitophagy, impairing a vital quality-control process that removes damaged mitochondria.

## Results

### Mfn1 and Mfn2 have opposite effects on PXA-induced mitochondrial matrix contractions

We previously described mitochondrial fragmentation and apoptosis caused by rapid mitochondrial Ca^2+^ release triggered by PXA (Bohler *et al*., 2018). To avoid downstream effects of mitochondrial fragmentation and apoptosis (Wang *et al*, 2025a), we began using Drp1 KO cells in further investigations of the mechanisms underlying PXA-induced Ca^2+^ release. Knockouts were confirmed by Western blotting (Fig. S1A). Mutations in Drp1 prevent apoptosis and mitochondrial outer membrane fission, but PXA still induces excessive Ca^2+^ release, leading to mitochondrial matrix contraction. Drp1 KO cells treated with PXA exhibit a “beads on a string” or “pearling” phenotype (Stepp *et al*, 2026), due to the localized constrictions resulting from these matrix contractions (Fig. 1A,B).

**Fig. 1.**
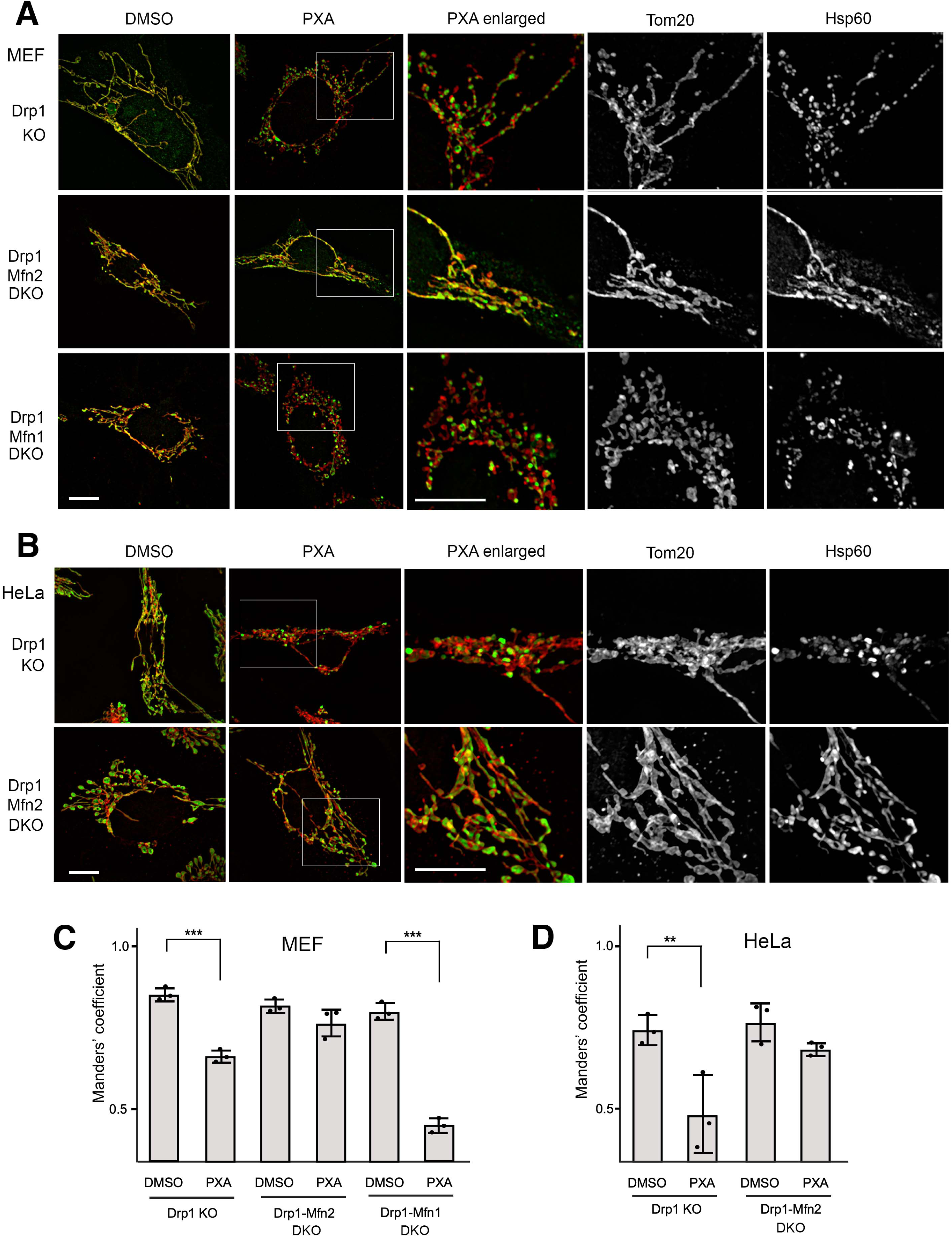
PXA-induced mitochondrial constrictions are controlled by Mitofusins. (**A**) Tom20 (red) and Hsp60 (green) immunofluorescence of MEFs with genotypes as indicated and 30 min treatments at 37 °C with DMSO or 10 μM PXA. Boxed areas were enlarged to show morphological changes. Scale bar is 10 μm. (**B**) Similar images with Drp1 KO and Drp1-Mfn2 DKO HeLa cells. (**C, D**) Manders’ coefficients of Tom20 and Hsp60 colocalization to quantify matrix contraction. Averages ± SD from three independent experiments (50 images per condition) analyzed by one-way ANOVA with Tukey’s HSD post hoc test.

Observing localized constrictions induced by PXA in Drp1 KO cells prompted us to investigate possible roles of other mitochondrial dynamics proteins in this process. The roles of Mfn1 and Mfn2 were assessed with Mfn1-Drp1 and Mfn2-Drp1 double-knockout (DKO) MEFs, as well as in Drp1-Mfn2 DKO HeLa cells, where Drp1 KO served to prevent apoptosis. Loss of each protein was confirmed by Western blotting (Fig. S1A). Strikingly, PXA elicited markedly different responses in Drp1-Mfn1 versus Drp1-Mfn2 DKO cells. Drp1-Mfn2 DKO MEFs and HeLa cells were completely resistant to PXA-induced matrix contraction, whereas Drp1-Mfn1 DKO cells exhibited enhanced matrix contraction compared with Drp1 knockout cells. In fact, this contraction was sufficient to drive mitochondrial fission even in the absence of Drp1 (Fig. 1A,B).

PXA-induced matrix retraction in Drp1 KO cells causes separation between outer-membrane and matrix fluorescence signals, visible in scatterplots (Fig. S1B). This separation enables quantification using Manders’ coefficients to assess the overlap between an outer-membrane marker (Tom20) and a matrix marker (Hsp60). Manders’ coefficients showed a significant decrease in overlap in Drp1 KO cells treated with PXA, but not in Drp1-Mfn2 DKO cells. Conversely, the loss of overlap was more pronounced in Drp1–Mfn1 DKO cells (Fig. 1C,D). These findings suggest that Mfn2 inhibits PXA-induced contraction, while Mfn1 promotes it. This functional opposition aligns with previous siRNA studies in cardiomyocytes that demonstrated Mfn1 and Mfn2 differentially affect mitochondrial Ca²⁺ uptake and release (Inagaki *et al*, 2023). Their opposing roles may arise from both their distinct functions and their ability to heterodimerize and counteract one another. In summary, Mfn2 is essential for PXA-induced mitochondrial matrix contraction and fission, whereas Mfn1 acts as an antagonist to this process.

### Mfn2 controls mitochondrial Ca^2+^ release through NCLX when induced by PXA

We focused on NCLX-mediated Ca^2+^ release because Mfn2 was previously shown to interact with and affect NCLX (Samanta *et al*., 2018). PXA-induced mitochondrial Ca^2+^ release was monitored in HeLa cells using a mitochondria-targeted fluorescent Ca^2+^ reporter (R-Cepia3mt) and the fluorescent Ca^2+^ dye Rhod-2 AM. HeLa cells were used throughout for Ca^2+^ release experiments because those cells yield good signal-to-noise ratios with the available probes. Single-cell Ca^2+^ traces were collected with the R-Cepia3mt fluorescent protein using spinning-disk confocal microscopy (Fig. 2A,B), and population-level responses were measured with Rhod-2 AM using a plate reader (Fig. S2B). A CRISPR/Cas9-generated NCLX-KO HeLa line, validated by western blotting and comparison with NCLX-siRNA samples (Fig. S2A), was used as a control. Consistent with prior observations in WT cells (Bohler *et al*., 2018), PXA caused rapid mitochondrial Ca^2+^ release in Drp1-KO cells (Fig. 2A,B, Fig. S2B-D). In contrast, this Ca^2+^ release was absent in Mfn2-Drp1 DKO cells, NCLX-KO cells, Drp1- NCLX DKO cells, and Drp1 KO cells treated with the NCLX inhibitor CGP37157 (Fig. 2A,B). These findings demonstrate that PXA-induced mitochondrial Ca^2+^ release depends on both Mfn2 and NCLX.

**Fig. 2.**
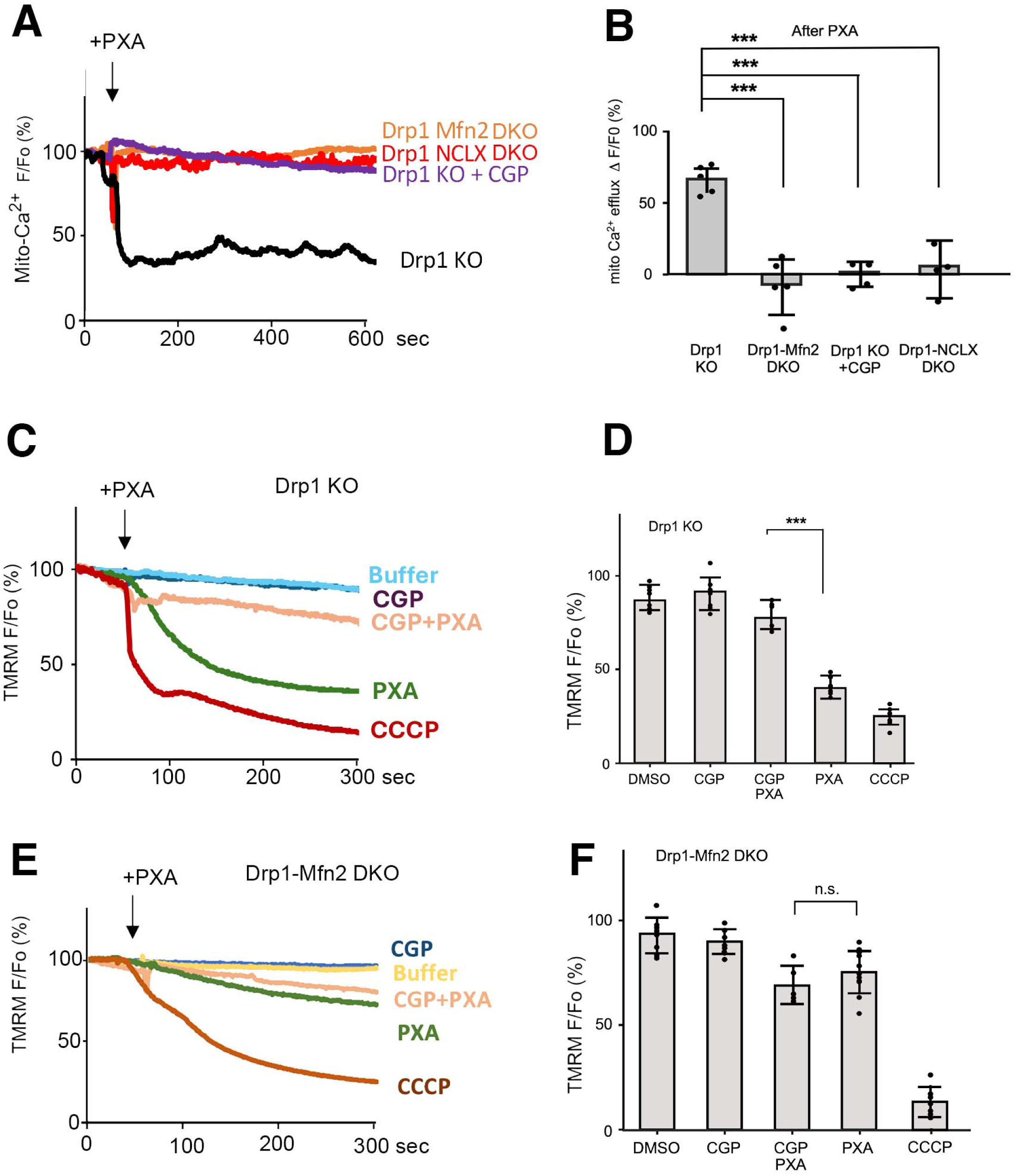
Mfn2 promotes mitochondrial Ca^2+^ efflux and membrane potential through NCLX. (**A**) Tracings of mitochondrial matrix Ca^2+^ detected with matrix3mt-R-CEPIA in HeLa cells with genotypes as indicated. Baseline (Fo) was established using 60-sec tracings. Ca^2+^ release was induced with 10 μM PXA, followed by 600 sec of further recording. Where indicated, cells were pre-incubated for 30 min at 37 °C with 10 μM CGP37157. (**B**) Relative fluorescence (F/Fo) at 600 sec, reflecting changes in mitochondrial Ca^2+^ levels. (**C**) Effects of PXA on mitochondrial membrane potential (ΔΨ) in Drp1 KO HeLa cells stained with 25 nM TMRM. Where indicated, cells were preincubated for 30 min with 10 μM CGP37157 (NCLX inhibitor). Baselines and tracings after perfusion with or without 10 μM PXA were obtained as in panel A. Perfusion with 10 μM CCCP was used as a control for dissipation of ΔΨ. (**D**) Averages of F/Fo were determined at 300 seconds after perfusion. (**E,F**) Tracings and histogram of TMRM fluorescence with Drp1-Mfn2 DKO cells instead of Drp1 KO cells. Throughout, averages with SD are shown with significance analyzed by one-way ANOVA with Tukey’s HSD post hoc test.

We then assessed mitochondrial membrane potential, which was previously shown to decrease after PXA treatment, likely as a downstream effect of Ca^2+^ release. Ca^2+^ export by NCLX is electrogenic: each Ca^2+^ ion is exchanged for three Na^+^, which reduces the membrane potential, as this was recently shown to have a large Na^+^ ion component (Hernansanz-Agustin, 2025). The resulting mitochondrial Na^+^ load is subsequently exchanged for protons via the mitochondrial Na^+^/H^+^ exchanger (Hernansanz-Agustin *et al*, 2024; Numata *et al*, 1998). This proton influx reduces the proton gradient and reduces matrix pH without further changes in membrane potential. Our data show that PXA-induced membrane depolarization is prevented by Mfn2 loss and by CGP37157 (Fig. 2C-F). Therefore, the loss of mitochondrial membrane potential is an indirect result of NCLX activation by PXA. PXA-induced Ca^2+^ release does not occur in Mfn2-Drp1 DKO cells, indicating that it is specific to the role of Mfn2 in promoting NCLX activity. In contrast, treatment with CCCP causes Ca^2+^ release in all tested genotypes (Fig. S2E-F), demonstrating that this response does not require Mfn2 or NCLX. Ca^2+^ release induced by CCCP is likely induced by changes in the matrix pH, which can solubilize matrix-buffered Ca^2+^ (De Stefani *et al*, 2016), while release is mediated by MCU in mitochondria lacking membrane potential (Montero *et al*, 2001). Unlike the effects of CCCP, we find that loss of membrane potential induced by PXA results from Na^+^/Ca^2+^ exchange.

### PXA interacts directly with Mfn2 in a complex with NCLX

We used a cellular thermal shift assay (CETSA) to determine if PXA directly binds to Mfn2 or NCLX. In this assay, MEF extracts, solubilized with dodecyl maltoside, were incubated with PXA and subjected to a temperature gradient. After heat-induced denaturation, insoluble proteins were removed by centrifugation, and the remaining soluble proteins were analyzed by western blots. To evaluate shifts in denaturation temperature, we examined MEFs overexpressing Mfn2-FLAG and MEFs lacking Mfn1 or Mfn2. The functionality of Mfn2-FLAG was confirmed by its ability to rescue the mitochondrial morphology defect in Mfn2-KO cells (Fig. S3A). Since available NCLX antibodies recognize multiple nonspecific bands, we verified the identity of the NCLX band using KO controls (Fig. S3B, S3C). The CETSA results indicate that PXA shifts the denaturation temperatures of both Mfn1 and Mfn2, but not NCLX or SLC25A46, a mitochondrial outer membrane protein associated with Mitofusins and Opa1 (Steffen *et al*, 2017) (Fig. 3A,B). Vinculin, used as a control, remained unaffected. The thermal stabilization by PXA was more pronounced in cells overexpressing Mfn2, supporting a direct interaction between PXA and Mfn2. Although PXA also influences Mfn1, suggesting it binds to additional proteins or conserved regions shared by Mfn1 and Mfn2, this does not change the essential role of Mfn2 in PXA-induced Ca^2+^ release. Several other targets of PXA were identified (Ali *et al*, 2022; Ceccacci *et al*, 2020), but the concentrations used in those studies were much higher than in our experiments, suggesting that they are not the primary cause of the morphological defects induced by PXA.

**Fig. 3.**
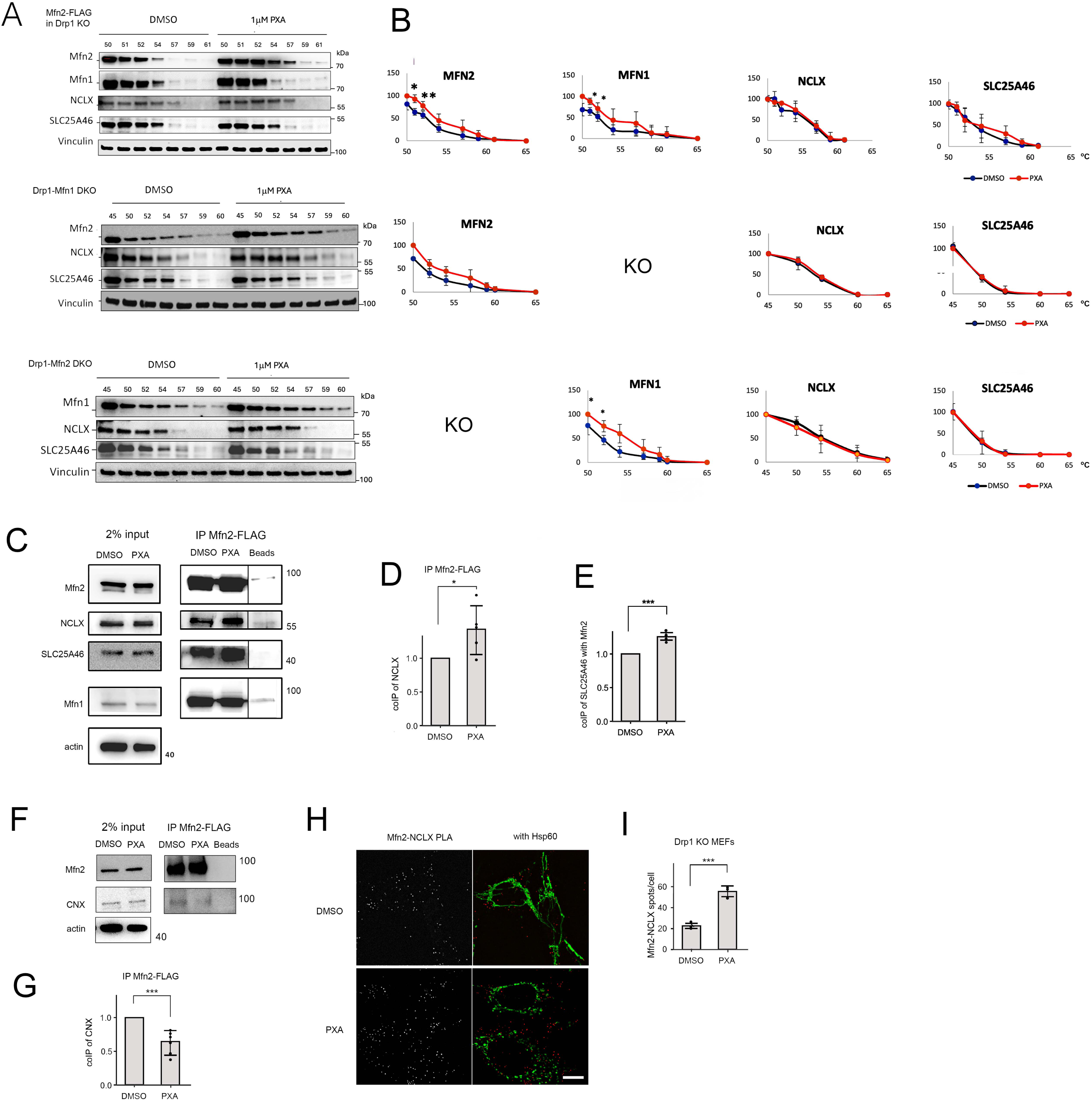
PXA targets Mitofusins and induces interactions between Mfn2 and NCLX. (**A**) CETSA to determine the target of PXA. Dodecyl maltoside extracts from cells with the indicated genotypes were incubated with DMSO or 1μM PXA and subjected to heat denaturation for 3 min at the indicated temperatures, followed by removal of denatured proteins by centrifugation and western blot analysis of the supernatants. The upper panel was made with Drp1-Mfn2 DKO cells that stably express exogenous Mfn2-FLAG, while the lower two sets of blots were made with endogenous proteins. (**B**) Band intensities determined with densitometric scans of blots as shown in panel A. The intensities were normalized to the bands at 45 or 50°C, showing averages with SD from 3 independent experiments. (**C**) Co-immunoprecipitation (coIP) of endogenous NCLX and SLC25A46 with Mfn2-FLAG using FLAG antibody attached to beads. Cells grown under glycolytic conditions (Gly) were treated for 30 min with DMSO, 10 μM PXA. These cells were incubated with DSP cross-linker and subjected to coIP, followed by western blot analysis. Actin was used as a loading control. (**D**) Densitometry of the NCLX coIPs, normalized to the levels of Mfn2 for each condition. Averages of 5 independent experiments are shown with SD and results of an unpaired Student’s t-test. (**E**) Densitometry of SLC25A46 coIPs as described for NCLX in panel D. Averages of 3 independent experiments with SD and results of an unpaired Student’s t-test are shown. (**F**) CoIP of Calnexin with Mfn2-FLAG after treatment with PXA under glycolytic conditions as described in panel C. (**G**) Densitometry of Calnexin coIP with Mfn2-FLAG as in panels D and E. Averages of 3 independent experiments with SD and results of an unpaired Student’s t-test are shown. (**H**) PLA of Mfn2-myc and NCLX-HA (red spots) in glycolytic Drp1 KO MEFs treated for 30 min with DMSO or 10 μM PXA. Mitochondria were detected by immunofluorescence with a chicken antibody against Hsp60 (green). The size marker is 10 μm. (**I**) Average numbers of PLA spots per cell from 3 independent experiments with SD and results of an unpaired Student’s t-test are shown. PLA spots in 35 cells were counted for each experiment.

To determine whether Mfn2 directly interacts with NCLX, we conducted co-immunoprecipitation assays using epitope-tagged Mfn2-FLAG and chemical cross-linking. Mfn2-myc was confirmed to be functional by its ability to rescue the mitochondrial fusion defect in Mfn2 KO cells (Fig. S3A). Western blots showed increased co-immunoprecipitation of NCLX with Mfn2 after PXA treatment, along with a higher association of SLC25A46 (Fig. 3C-E). Conversely, co-immunoprecipitation of Calnexin decreased following PXA treatment, indicating that Mfn2 dissociates from the MAM under these conditions (Fig. 3F,G).

We further examined Mfn2-NCLX proximity using a Proximity Ligation Assay (PLA). Cells were transfected with Mfn2-myc and NCLX-HA, cultured under glycolytic conditions, and processed for PLA with myc and HA antibodies. Mitochondria were labeled using a chicken anti-Hsp60 antibody. PLA analysis showed an increased number of PLA puncta in glycolytic cells treated with PXA (Fig. 3H,I). PLA spots did not always coincide with Hsp60 staining because the mitochondrial matrix appears more condensed than the membranes where Mfn2 and NCLX are located. Co-staining with the outer membrane marker TOM20 confirmed that PLA signals originate from mitochondria (Fig. S3D). These findings demonstrate that Mfn2 and NCLX are in closer proximity after treatment with PXA. Overall, the results suggest that Mfn2 is essential for mitochondrial Ca²⁺ release via NCLX, and that this function is activated by PXA, which directly targets Mfn2.

### Mfn2 controls ROS-induced mitochondrial Ca^2+^ release through NCLX

Mitochondrial Ca^2+^ release is often monitored following extracellular histamine or ATP application, but these methods focus on the homeostatic functions of NCLX at the MAM. These methods also make it difficult to distinguish the tethering functions of Mfn2 at the MAM from a potential role in promoting NCLX activity without a detailed understanding of both functions. Instead, we assessed the physiological role of Mfn2-regulated Ca^2+^ release by NCLX under conditions that increase mitochondrial ROS. These conditions induce mitochondrial damage and, as a result, activate mitophagy to remove dysfunctional mitochondrial parts. Common methods for inducing mitophagy, such as CCCP or oligomycin/antimycin A treatments in glycolytic cells (Dong *et al*, 2023), were deemed uninformative because loss of mitochondrial membrane potential also prevents electrogenic Ca^2+^ release by NCLX. To maintain mitochondrial membrane potential, we induced ROS by treating respiring cells (grown with galactose) with oligomycin or mitoPQ. Oligomycin, an ATP synthase inhibitor, causes ATP depletion but also produces ROS in respiring cells through hyperpolarization of the mitochondrial inner membrane (Wilkins *et al*, 2022), while mitoPQ is a mitochondrial-targeted derivative of paraquat that uses electrons from the ETC to generate ROS (Antonucci *et al*, 2019).

We used N-acetyl cysteine (NAC) treatments to determine whether the effects on Ca^2+^ release and the Mfn2-NCLX complex are due to increased ROS levels. Cells were pre-incubated for 30 minutes with 5 mM NAC before inducing Ca^2+^ release with oligomycin and mitoPQ in WT HeLa cells grown under respiring conditions. As a control, we detected ROS with the ROS-sensitive dye mitoSOX. As expected, ROS levels increased with mitoPQ and oligomycin treatments and were reduced by pre-treatment with NAC (Fig. S4A). We found that oligomycin and mitoPQ treatments under respiring conditions (growth with galactose) induce mitochondrial Ca^2+^ release in HeLa cells, and this release can be blocked by quenching ROS with N-acetyl cysteine (NAC) (Fig. S4B, C), showing that ROS are needed for this response. Oligomycin does not decrease mitochondrial membrane potential (Wilkins *et al*., 2022), which could have been a factor, because Ca^2+^ release by NCLX also depends on mitochondrial membrane potential (Islam *et al*, 2020). However, we also tested whether mitoPQ affects membrane potential. Our results show that 50 and 100 μM mitoPQ do not impact mitochondrial membrane potential (Fig. S4D, E), thus supporting ROS-induced Ca^2+^ release.

We used gene knockout to examine how Mfn2 and NCLX affect mitochondrial Ca^2+^ release triggered by oligomycin and mitoPQ. Knockouts of Mfn2 and NCLX were created in a Drp1 KO background to avoid complications from mitochondrial fragmentation during the assay. Our results show that Mfn2-Drp1 and NCLX-Drp1 DKO cells have impaired Ca^2+^ release, similar to what is seen with CGP37157 treatment (Fig. 4A-D). To confirm that these effects were due to specific gene knockdowns, we also tested siRNA effects. Successful knockdown of Mfn2 and NCLX in WT HeLa cells was confirmed by western blot (Fig. S4F). Similar to gene knockouts, Ca^2+^ release is also blocked by NCLX or MFN2 siRNA and by the NCLX inhibitor CGP37157 (Fig. S4G-J). The reduced mitochondrial Ca^2+^ observed suggests that Ca^2+^ is either released into the cytosol or transferred to other organelles, such as the ER or lysosomes. We tested this with the cytosolic Ca^2+^ dye Calbryte-520 AM. Drp1 KO HeLa cells treated with oligomycin or mitoPQ showed a strong, sustained increase in cytosolic Ca^2+^ (Fig. 4E-H). This increase was prevented in Mfn2-Drp1 and NCLX-Drp1 DKO cells, as well as in Drp1 KO cells treated with CGP37157. We conclude that oligomycin and mitoPQ induce Ca^2+^ release into the cytosol of respiring HeLa cells, and this process requires both Mfn2 and NCLX.

**Fig. 4.**
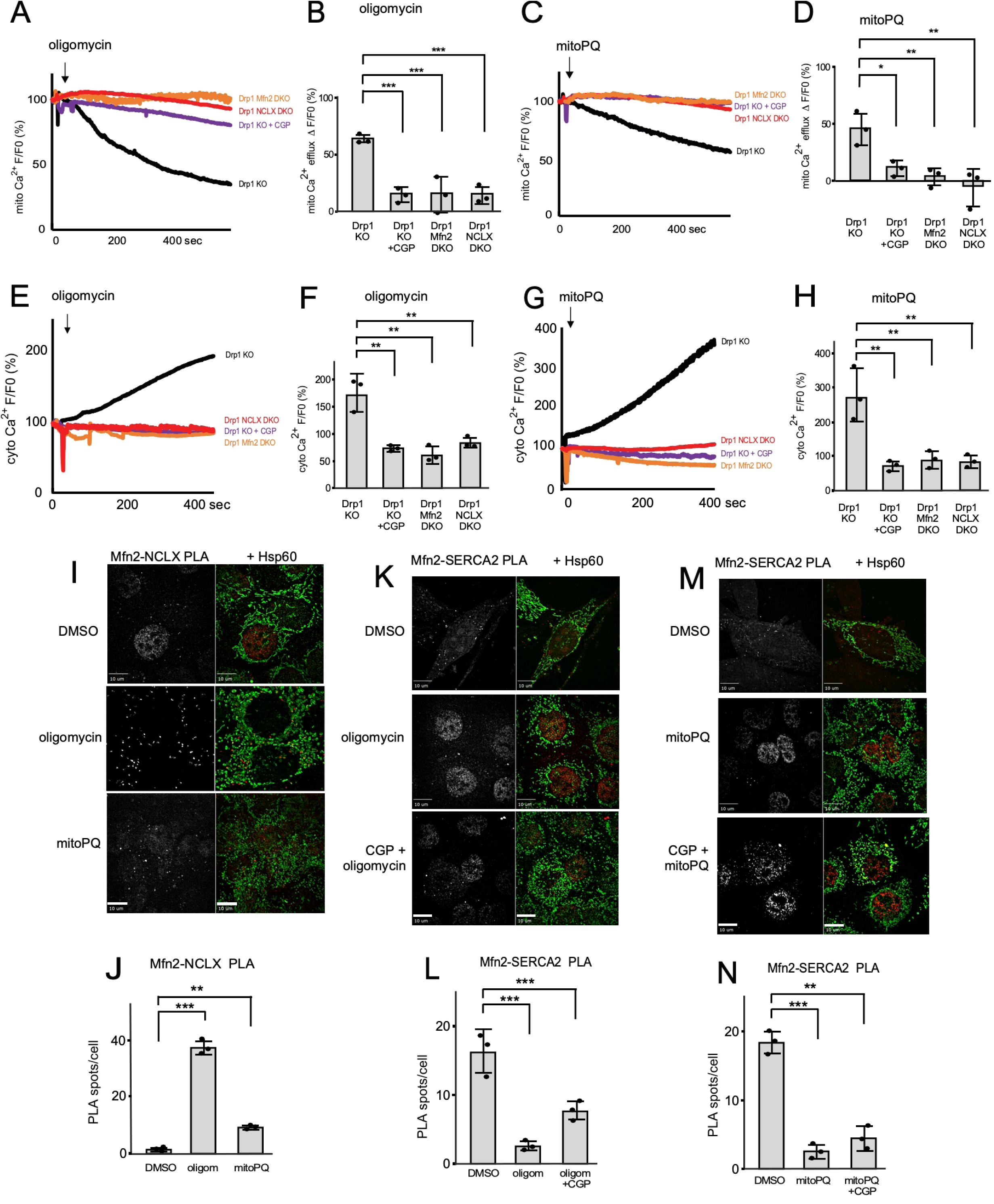
Stress-induced mitochondrial Ca^2+^ release is mediated by NCLX in association with Mfn2. (**A**) Tracings of mitochondrial Ca^2+^ fluorescence detected with matrix-3mt-CEPIA in HeLa cells with genotypes as indicated. Measurements were as described in Fig. 2A. Where indicated, cells were pre-incubated for 30 min at 37 °C with 10 μM CGP37157. (**B**) Quantification of mitochondrial Ca^2+^ levels, expressed as F/Fo at 400 sec after oligomycin treatment, as in Fig. 2B.(**C, D**) Tracings and histograms of mitochondrial matrix Ca^2+^, as in panels A and B but treated with 50 μM mitoPQ. (**E**) Tracings of cytosolic Ca^2+^ induced in respiring Hela cells with 10 μM oligomycin and detected with Calbryte-520 AM. Where indicated, cells were pre-incubated for 30 min with 10 μM CGP37157. (**F**) Changes in cytosolic Ca^2+^ levels reflected by the relative fluorescence (F/Fo) at 600 sec. (**G,H**) Tracings and histograms of cytosolic Ca^2+^ levels as in panels E and F but induced with 50 μM mitoPQ. (**I**) PLA of Mfn2-myc and NCLX-HA (red spots) in respiring Drp1 KO MEFs treated with DMSO, 10 μM oligomycin, or 50 μM mitoPQ. Mitochondria were detected by immunofluorescence with chicken Hsp60 antibody (green). (**J**) Number of PLA spots per cell, averaged across 25 cells for each condition in three rounds of experiments. (**K**) PLA of Mfn2-myc and endogenous SERCA2 (red spots) in respiring Drp1 KO MEFs treated and quantified as in panel J. (**L**) Numbers of PLA spots per cell analyzed as described in panel J. (**M,N**) PLA of Mfn2-myc and endogenous SERCA2 (red spots) in respiring Drp1 KO MEFs treated and quantified as in panel J. The size markers in panels I, K and M are 10 μm. Throughout, averages are shown with SD and significance was determined with one-Way ANOVA with Tukey’s HSD post hoc test.

To determine whether interactions between Mfn2 and NCLX are enhanced under ROS-inducing conditions, we performed PLA with Mfn2-myc and NCLX-HA and co-stained for mitochondria using chicken anti-Hsp60. We observed an increase in Mfn2/NCLX PLA puncta in cells treated with oligomycin or mitoPQ (Fig. 4I,J), indicating closer proximity between Mfn2 and NCLX under these conditions. We pretreated with NAC to test whether the association between Mfn2 and NCLX detected by PLA is promoted by ROS under these conditions. PLA for Mfn2 and NCLX, with or without pre-incubation with NAC, shows that NAC suppresses the association between Mfn2 and NCLX (Fig. S4K,L), which aligns with NAC’s inhibitory effects on Ca^2+^ release. Having confirmed that Mfn2 associates more strongly with NCLX during stress and that Mfn2 is necessary for NCLX-dependent mitochondrial Ca^2+^ release, we then examined whether Mfn2’s association with the MAM is similarly affected, using PLA for SERCA2 and transiently expressed Mfn2-myc in Drp1 KO MEFs. Treatment with oligomycin and mitoPQ, or in combination with the NCLX inhibitor CGP37157, each led to a significant decrease in Mfn2-SERCA2 PLA spots (Fig. 4K-N), suggesting reduced Mfn2 interaction with ER-mitochondria contact sites. These results indicate that oligomycin and mitoPQ increase Mfn2’s association with NCLX, while they decrease Mfn2’s association with SERCA2 at ER-mitochondrial contact sites.

### SLC25A46 promotes Ca^2+^ release and the association between Mfn2 and NCLX

To understand how Mfn2, which is anchored in the mitochondrial outer membrane with only a small part exposed to the intermembrane space, can assist Ca^2+^ release by NCLX on the mitochondrial inner membrane, we examined several Mfn2-interacting proteins that have larger exposed regions in the intermembrane space as potential intermediaries between Mfn2 and NCLX. These proteins include Mtch1, Mtch2, MarchV, and SLC25A46. We focused on SLC25A46 because previous research showed that the mitochondrial fusion proteins (Mfn1, Mfn2, and Opa1) co-immunoprecipitate with this protein (Steffen *et al*., 2017), and it plays a role in stress-induced mitochondrial dynamics (Schuettpelz *et al*, 2023).

Mitochondrial Ca^2+^ dynamics were observed in WT HeLa cells and cells transfected with SLC25A46 siRNA. Successful SLC25A46 knockdown was confirmed by Western blotting (Fig. S5A). Mitochondrial Ca^2+^ traces indicate that SLC25A46 knockdown decreases mitochondrial Ca^2+^ efflux triggered by oligomycin or mitoPQ under respiring conditions (Fig. 5A,B). To determine whether SLC25A46 is necessary for the interaction between Mfn2 and NCLX, we compared PLA signals for Mfn2-myc and NCLX-HA in Drp1 KO MEFs and SLC25A46-Drp1 DKO cells, with or without oligomycin and mitoPQ treatments. The knockouts were verified by Western blot (Fig. S5B). The results show a nearly complete loss of Mfn2-NCLX spots when SLC25A46 is knocked out (Fig. 5C,D). These effects were confirmed with coIP of NCLX with Mfn2-FLAG in WT but not in SLC25A46 KO HeLa cells (Fig. 5E). SLC25A46 KO in HeLa cells and comparable levels of Mfn2-FLAG expression in stable cell lines were validated with western blots (Fig. S5C,D). We conclude that SLC25A46 is essential for interactions between Mfn2 and NCLX, as well as for ROS-induced Ca^2+^ release.

**Fig. 5.**
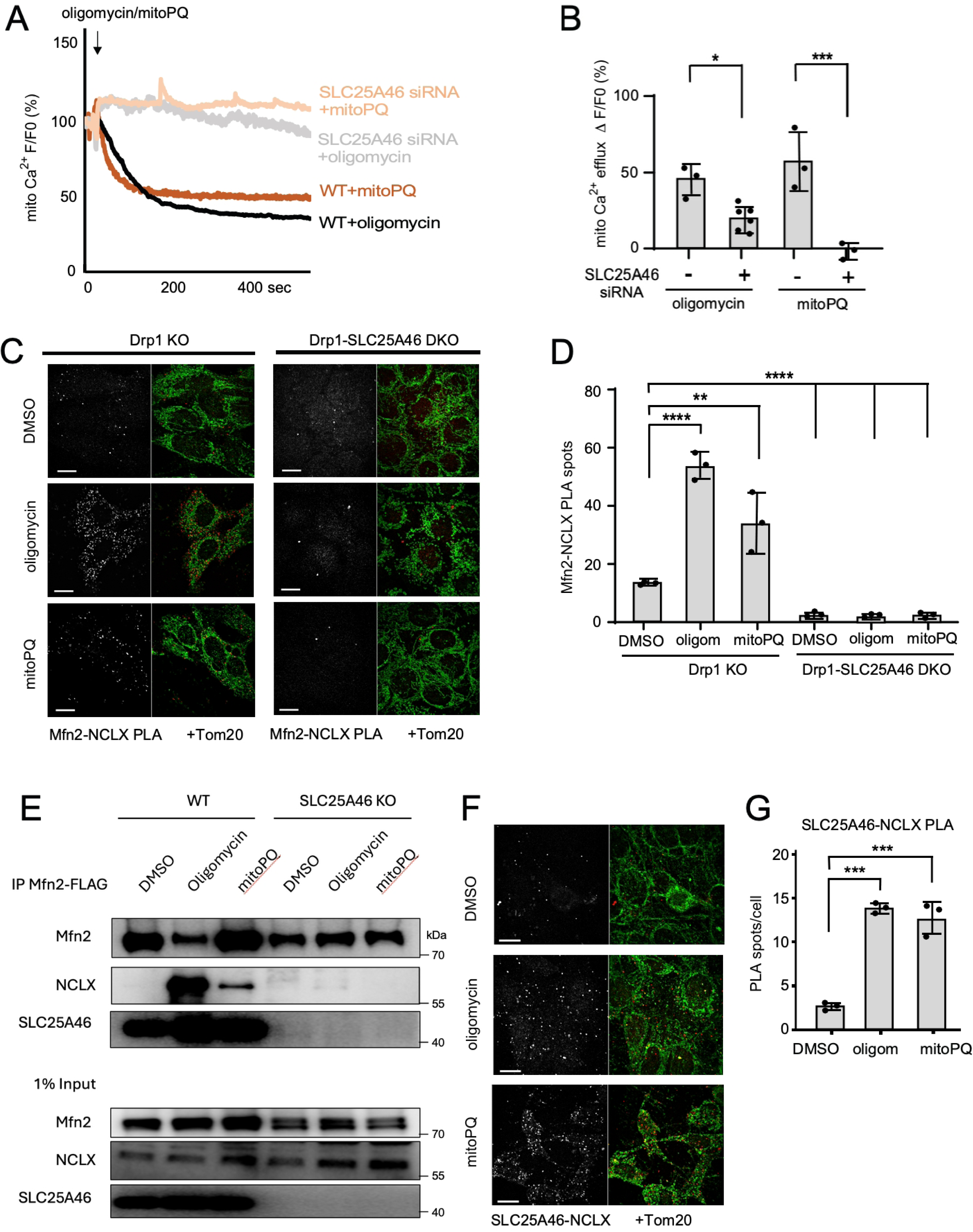
SLC25A46 is required for Mfn2-NCLX interactions and stress-induced Ca^2+^ release. (**A**) Tracings of mitochondrial matrix Ca^2+^ in respiring HeLa cells, detected as in Fig. 2A, but induced with 10 μM oligomycin or 50 μM mitoPQ, and in cells that were transfected with siRNA for SLC25A46. (**B**) Changes in mitochondrial Ca^2+^ levels reflected by the relative fluorescence (F/Fo) at 600 sec. (**C**) PLA of Mfn2-myc and NCLX-HA (red spots) in respiring Drp1 KO or SLC25A46 Drp1 KO MEFs treated with DMSO or 10 μM oligomycin, or 50 μM mitoPQ. Mitochondria were detected with chicken Tom20 antibody (green). The size marker is 10 μm. (**D**) Number of PLA spots for Mfn2-myc and NCLX-HA per cell, averaged across 25 cells for each condition in three rounds of experiments. (**E**) Western blots showing coIP of NCLX and SLC25A46 with Mfn2-FLAG in WT and SLC25A45 KO cells. (**F**) PLA of SLC25A46-myc and NCLX-HA (red spots) in respiring Drp1 KO MEFs treated for 30 min with DMSO, 10 μM oligomycin, or 50 μM mitoPQ. Mitochondria were detected with chicken Tom20 antibody (green). The size marker is 10 μm. (**G**) Number of PLA spots for SLC25A46-myc and NCLX-HA per cell, averaged across 25 cells for each condition in three rounds of experiments.

Next, we examined interactions between SLC25A46 and NCLX and between Mfn2 and SLC25A46 to determine which pairs change under ROS-induced conditions. We observed more PLA spots with SLC25A46-myc and NCLX-HA in Drp1 KO cells treated with oligomycin or mitoPQ (Fig. 5F,G), along with increased coIP of NCLX-HA with SLC25A46-myc (Fig. S5E,F), but no increase in coIP of Mfn2 with SLC25A46 or vice versa of SLC25A46 with Mfn2 (Fig. S5G,H). These results indicate that the interactions between Mfn2 and SLC25A46 are stable, while those with NCLX depend on SLC25A46 and are induced by ROS.

### Alternative pathways for control of mitochondrial Ca^2+^ release and fission

Opa1 is a potential regulator of Mfn2-NCLX interactions, since Opa1 also interacts with SLC25A46 (Schuettpelz *et al*., 2023; Steffen *et al*., 2017). Opa1 is controlled by Oma1-mediated proteolytic cleavage (Ehses *et al*, 2009; Head *et al*, 2009), which is triggered by oligomycin in respiring cells (MacVicar & Lane, 2014), as also observed by us (Fig. S6A). To determine whether Oma1 influences Mfn2-NCLX interactions and Ca^2+^ release, we examined the effects in Oma1 KO HeLa cells (KO confirmed by Western blot, Fig. S6B) with or without oligomycin in respiring cells. We found that Oma1 KO prevented oligomycin-induced mitochondrial fragmentation in respiring cells, as shown by immunofluorescence images and aspect ratios (Fig. 6A, B). This lack of fission is like that observed in glycolytic Oma1 knockout cells treated with CCCP (Quiros *et al*, 2012). Moreover, PLA results indicate that the oligomycin-induced increase in Mfn2-NCLX interactions is also blocked in Oma1 KO cells (Fig. 6C, D).

**Fig. 6.**
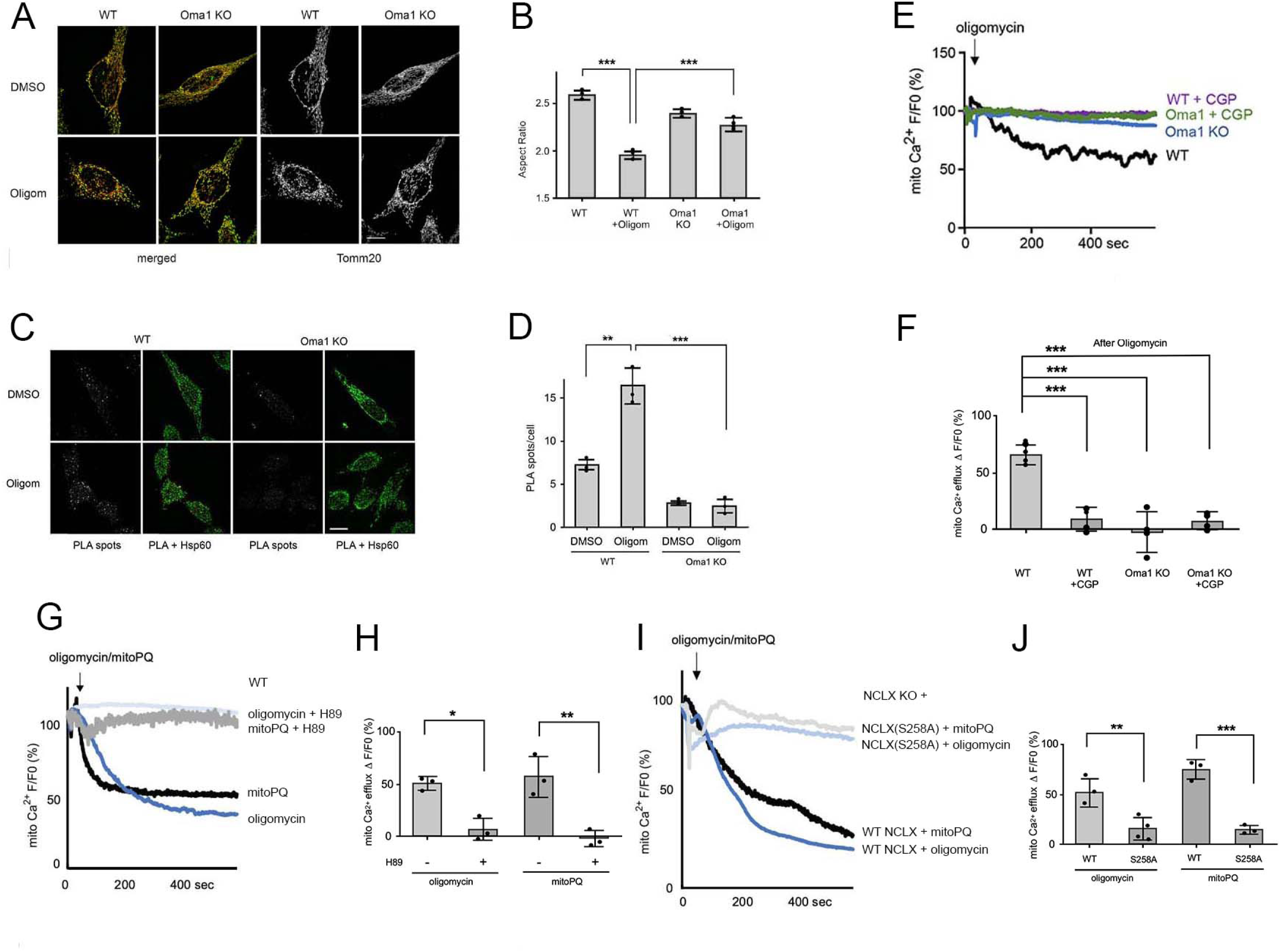
Mitochondrial ROS activation of Oma1 and PKA triggers Ca^2+^ release. (**A**) Immunofluorescence of WT and Oma1 KO HeLa cells treated for 30 min with DMSO or 10 μM oligomycin in respiring conditions. Merged images show Tomm20 (red) and Hsp60 (green), while the black and white images show Tomm20 labeling separately. Scale bar is 10 μm. (**B**) Aspect ratios of mitochondrial outer membranes (Tom20) under the conditions for panel B. Aspect ratios were determined for each condition with mitochondria in 25 cells per experiment in 3 independent experiments with SD and results of a one-way ANOVA. (**C**) PLA of Mfn2-myc and NCLX-HA (red spots) in WT and Oma1 KO HeLa cells grown under respiring conditions and treated for 30 min with DMSO or 10 μM Oligomycin. Mitochondria were detected by immunofluorescence with chicken Hsp60 antibody (green). Scale bar is 10 μm. (**D**) Numbers of PLA spots per cells treated with DMSO or oligomycin from 3 independent experiments with SD and results of a one-way ANOVA. (**E**) Tracings of mitochondrial matrix Ca^2+^, detected as in Fig. 2A, in WT and Oma1 KO HeLa cells grown under respiring conditions and treated with 10 μM oligomycin. Where indicated with CGP, cells were preincubated for 30 min at 37 °C with 10 μM CGP37157. (**F**) Changes in mitochondrial Ca^2+^ levels are reflected by the relative fluorescence (F/Fo) at 600 sec. Averages are shown with SD, and significance is determined with one-way ANOVA. (**G**) Tracings of mitochondrial matrix Ca^2+^ in respiring HeLa cells, detected as in Fig. 2A, but induced with 10 μM oligomycin or 50 μM mitoPQ, with or without pretreatment for 30 min with 10 μM H89 (PKA inhibitor). (**H**) Relative fluorescence (F/Fo) at 600 sec, reflecting changes in mitochondrial Ca^2+^ levels. (**I**) Tracings of mitochondrial matrix Ca^2+^ in respiring HeLa cells, detected as in Fig. 2A, but induced with 10 μM oligomycin or 50 μM mitoPQ, in NCLX KO cells that stably express exogenously introduced wt NCLX-HA or NCLX(S258A)-HA. (**J**) Quantification of mitochondrial Ca^2+^ efflux rates of F/Fo. Averages are shown with SD and significance was determined with one-way ANOVA with Tukey’s HSD post hoc test.

To confirm that Oma1 effects result from Opa1 loss due to cleavage, we tested the complete absence of Opa1 using Drp1-Opa1 DKO cells (DKO confirmed by Western blotting, Fig. S6C). These cells showed more PLA spots than Drp1 KO cells, even without oligomycin treatment, and no further increase after oligomycin was applied (Fig. S6D, E). Mitochondrial Ca^2+^ release was blocked in Oma1 KO cells, like the effects seen with the NCLX inhibitor CGP37157 (Fig. 6E, F). We conclude that Oma1-mediated cleavage of Opa1 promotes oligomycin-induced mitochondrial fission and the interaction between Mfn2 and NCLX, which is essential for Ca^2+^ release. Surprisingly, mitoPQ did not cause Opa1 cleavage in all cell types tested even though it still triggers Ca^2+^ release. These findings suggest that removing L-Opa1 during mitochondrial Ca^2+^ release is not always necessary. Oligomycin has been shown to induce Opa1 cleavage in respiring cells (MacVicar & Lane, 2014), and it may be more effective than mitopPQ, because ATP deficiency by itself is a strong inducer of Opa1 cleavage (Baricault *et al*, 2007; Rainbolt *et al*, 2016). Further research is needed to understand how L-Opa1 influences the interactions between Mfn2 and NCLX.

It was previously shown that NCLX is also activated by PKA-mediated phosphorylation at Ser258 (Kostic *et al*., 2015). We tested whether PKA is necessary for Ca^2+^ release induced by oligomycin and mitoPQ using the PKA inhibitor H89. The results demonstrate a block in Ca^2+^ release, indicating that PKA is essential for the ROS-induced activation of NCLX (Fig. 6G, H). To investigate this further, we expressed wild-type and phosphorylation-defective NCLX(S258A) in NCLX KO cells and treated them with oligomycin and mitoPQ. Expression levels of exogenously introduced NCLX were verified with Western blots (Fig. S6F). Cells expressing NCLX(S258A) showed impaired mitochondrial Ca^2+^ release compared to WT NCLX under ROS-inducing conditions (Fig. 6I, J). It was previously suggested that ROS can activate PKA via cysteine modification, thus bypassing cAMP-dependent activation (Ekhator *et al*, 2025; Zhang *et al*, 2016). It is therefore possible that ROS and PKA act in a linear pathway to promote mitochondrial Ca²⁺ release, although independent mechanisms of control cannot yet be ruled out.

To determine whether mitochondrial ROS, Ca^2+^ release, or lack of ATP, as sensed by AMPK, contribute to the mitochondrial fission observed with oligomycin and mitoPQ, cells were pre-incubated with NAC (a ROS scavenger), CGP37157 (an NCLX inhibitor), or BAY-3827 (an AMPK inhibitor) before treatment with oligomycin or mitoPQ. Fission was assessed using immunofluorescence with Tom20. Oligomycin caused significant mitochondrial fragmentation, while mitoPQ resulted in a milder phenotype. Pre-incubation with CGP37157 partially reduced fragmentation in mitoPQ-treated cells, but the effect on oligomycin-treated cells was not statistically significant. In contrast, NAC and BAY-3827 both prevented mitochondrial fission induced by oligomycin or mitoPQ (Fig. S6G,H). Since AMPK is activated by ATP depletion, these results suggest that stress-induced fission in these cells is mainly driven by ATP depletion rather than increased cytosolic Ca^2+^.

### Mitochondrial Ca^2+^ efflux promotes mitophagy through the NEDD4-1 E3 ubiquitin ligase

Instead of fission, Ca^2+^ release into the cytosol may influence other aspects of mitophagy. We tested this using the mitoQC reporter, a chimeric protein where GFP and mCherry are fused to the Fis1 tail anchor. Autophagolysosomes formed during mitophagy appear as red puncta because GFP is quenched in lysosomes. We found that oligomycin induces mitophagy in MEFs grown under respiring conditions, and this process is suppressed by NAC and CGP37157 (Fig. S7A,B). However, mitoPQ does not induce mitophagy in these cells, suggesting a more complex relationship between mitochondrial Ca^2+^ release and mitophagy. To explore whether cell-type-specific differences influence the ability of mitochondrial Ca^2+^ release to induce mitophagy, we examined its effects on mitophagy in SH-SY5Y cells. In these cells, both oligomycin and mitoPQ induce mitophagy, and this process is inhibited by NAC and CGP37157 (Fig. 7A-D). These findings suggest that SH-SY5Y cells, unlike MEFs, contain a Ca^2+^-sensitive factor that enables mitoPQ-induced mitophagy.

**Fig. 7.**
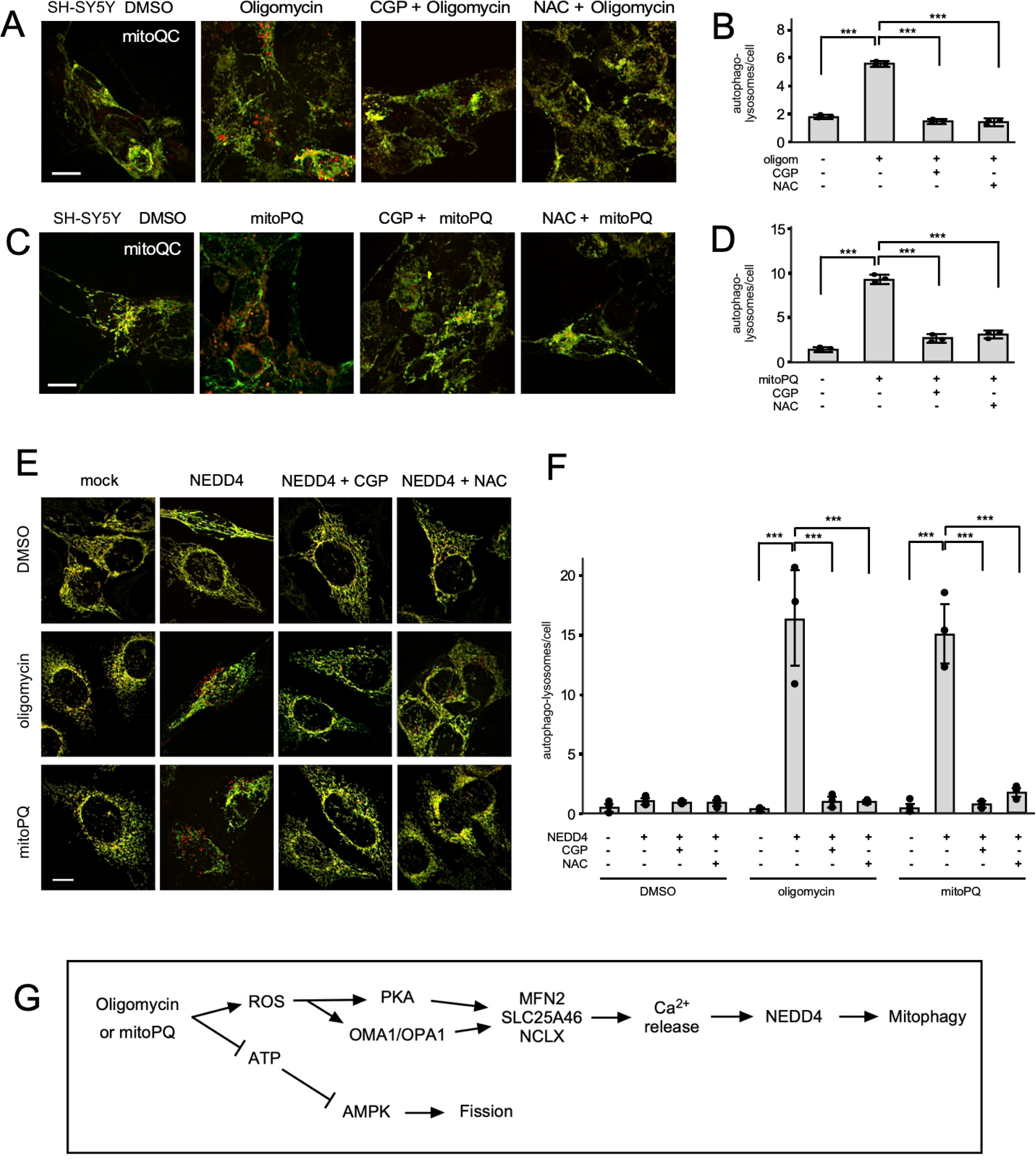
ROS-induced mitochondrial Ca^2+^ release promotes mitophagy by activating NEDD4-1. (**A**) Representative live images of WT SHY-5Y cells that stably express mitoQC (Fis1-mCherry-GFP) introduced with a lentivirus. Cells were pre-incubated with 10 μM CGP37157 or 5 mM NAC for 30 min at 37 °C and treated for 10 μM oligomycin for 30 min at 37 °C. Scale bar is 10 μm. (**B**) Quantification of red puncta per cell from 3 independent experiments with SD and results of one-Way ANOVA. (**C, D**) SHY-5Y cells were pre-incubated as in panels A and B, but then treated for 30 min with 50 μM mitoPQ. (**E**) HeLa cells that stably express mitoQC were transiently transfected with a Parkin construct (mock) or with Parkin and NEDD4-1 constructs (NEDD4), followed by preincubations with CGP37157 or NAC and treatments with oligomycin and mitoPQ as in panels A-D. (**F**) Quantification of red puncta per cell from 3 independent experiments with SD and results of one-Way ANOVA. (**G**) Proposed pathway linking ROS-induced mitochondrial Ca^2+^ release to cytosolic Ca^2+^ dependent activation of mitophagy.

To identify which mitophagy factors might detect mitochondrial Ca^2+^ release, we performed a targeted siRNA screen focusing on E3 ubiquitin ligases that could respond to cytosolic Ca^2+^. These include NEDD4-1 (Plant *et al*, 2000), NEDD4-L (Han *et al*, 2024), and SMURF1 (Lu *et al*, 2011), all of which have C2 domains, as well as TRIM5α (Saha *et al*, 2024)\ and UBE3 (Braganza *et al*, 2017). We found that NEDD4-1 siRNA reduces mitophagy induced by oligomycin and mitoPQ in SH-SY5Y cells, while siRNA against the other ligases has little or no effect (Fig. S7C-F). Knockdown was verified with Western blots (Fig. S7G). To verify that NEDD4-1 promotes Ca^2+^-induced mitophagy, we transfected HeLa cells with constructs expressing NEDD4-1 and Parkin (HeLa cells lack Parkin), or Parkin alone. Our results show that the combination of the two proteins, but not Parkin alone, significantly increases the number of autophagolysosomes formed in response to mitoPQ and oligomycin (Fig. 7E-F). Expression of NEDD4-1 was verified with Western blots (Fig. S7H). Parkin is likely required to initiate mitophagy when damaged mitochondrial fragments are generated, because these fragments lose membrane potential (Twig *et al*, 2008; Youle & van der Bliek, 2012). However, the role of NEDD4-1 in this process is novel. We conclude that NEDD4-1 promotes Ca^2+^-induced mitophagy in respiring cells. This effect is therefore dependent on NEDD4-1 expression levels, whereas in glycolytic HeLa cells, transfection with Parkin alone is sufficient to promote CCCP-induced mitophagy (Narendra *et al*, 2008).

Together, these results support a pathway in which ROS and PKA facilitate mitochondrial Ca^2+^ release by promoting interactions between NCLX, Mfn2, and SLC25A46 (Fig. 7G). Oma1-mediated cleavage of Opa1 strengthens these interactions while downstream effects of Ca^2+^ release on mitophagy through NEDD4-1 complement the effects of AMPK on mitochondrial fission and ULK1 activation during mitophagy (Iorio *et al*, 2021).

## Discussion

Mitochondria-derived ROS is vital for redox signaling and cellular stress responses, but a key question has remained unanswered: how are ROS produced in mitochondria converted into signals in the cytosol that indicate mitochondrial dysfunction? Here, we outline a mechanism that directly connects mitochondrial ROS to cytosolic signaling and mitophagy. We demonstrate that ROS causes NCLX-dependent Ca^2+^ release in a process that requires Mfn2, and that this Ca^2+^ signal promotes mitophagy in respiring cells. This pathway involves three coordinated components: (1) ROS-dependent signals that start Ca^2+^ release, (2) a protein complex spanning the mitochondrial inner and outer membranes that facilitates Ca^2+^ efflux, and (3) downstream Ca^2+^-dependent steps that trigger mitophagy. The newly identified role of Mfn2 in promoting mitochondrial Ca^2+^ release reveals a novel function, in addition to its well-known roles as a mitochondrial fusion protein (Chen *et al*., 2003) and as a tether linking mitochondria and the ER (de Brito & Scorrano, 2008).

Our data show that ROS stimulates mitochondrial Ca^2+^ release in respiring cells. As part of this process, ROS promotes the dissociation of Mfn2 from the MAM and increases the association between Mfn2 and NCLX. This increased association can be triggered by OMA1-dependent cleavage of OPA1 upon oligomycin treatment. In fact, the constant increase in Mfn2-NCLX interactions in Opa1 knockout cells suggests that Opa1 normally limits the formation of this complex. At the same time, PKA-dependent phosphorylation of NCLX enhances ROS-induced Ca^2+^ release, building on previous work on PKA regulation of NCLX activity (Kostic *et al*., 2015). Overall, these results highlight Mfn2 dissociation from the MAM as a key step in converting mitochondrial ROS into a cytosolic Ca^2+^ signal. Our data do not exclude the possibility that Mfn2 also supports basal NCLX-mediated Ca^2+^ efflux at the MAM, where SERCA2-dependent Ca^2+^ reuptake into the ER balances Ca^2+^ transfer from the ER to mitochondria through IP3R and MCU. However, the specific contribution of Mfn2 to NCLX regulation is difficult to separate from its established role as a tethering factor. Future studies employing artificial tethering systems will be required to directly address this possibility.

The ROS-driven Ca^2+^ release pathway described here originates within mitochondria and differs fundamentally from homeostatic fission triggered by external Ca^2+^ signals such as ER Ca^2+^ release or ionomycin-induced Ca^2+^ influx. Those pathways rely on INF2-mediated actin assembly at MAMs (Korobova *et al*, 2014; Korobova *et al*, 2013) and depend on mitochondrial Ca^2+^ uptake rather than Oma1 activation (Chakrabarti *et al*, 2018; Fung *et al*, 2023). While external signals such as PINK1- or PKA-dependent phosphorylation of Mfn2 may also promote ER dissociation and potentially influence Ca^2+^ release (Chen & Dorn, 2013; Dasgupta *et al*, 2021; Zhou *et al*, 2010), the pathway we define is initiated by mitochondrial ROS itself. Whether this ROS-driven mechanism occurs preferentially at mitochondrial tips and interfaces with lysosome-mediated fission processes (Kleele *et al*, 2021; Peng *et al*, 2020; Wong *et al*, 2019; Wong *et al*, 2018) remains to be determined. Nonetheless, these findings establish a new framework for understanding how mitochondria selectively identify and eliminate damaged regions through mitophagy (Twig *et al*., 2008; Youle & van der Bliek, 2012)

ROS-induced Ca^2+^ release is facilitated by a transmembrane complex involving Mfn2, NCLX, and the outer membrane protein SLC25A46. Although SLC25A46 interacts with both Mfn1 and Mfn2 (Steffen *et al*., 2017), their functions differ: Mfn1 works with L-OPA1 to promote stress-induced mitochondrial hyperfusion (Tondera *et al*., 2009), while Mfn2 serves as an ER–mitochondria tether (de Brito & Scorrano, 2008) and, as shown here, aids in mitochondrial Ca^2+^ release. Whether these differences come from distinct binding modes with SLC25A46 or involve additional cofactors remains unresolved. TMEM65, another inner membrane protein reported to associate with NCLX (Garbincius *et al*, 2025), has been suggested to regulate or even independently mediate Na^+^/Ca^2+^ exchange (Vetralla *et al*, 2025; Zhang *et al*, 2025), though this is still debated (Garbincius & Elrod, 2025). Our findings demonstrate that NCLX itself is essential for ROS-induced Ca^2+^ release, as NCLX-deficient cells do not show Ca^2+^ efflux in response to ROS, supporting a model in which TMEM65 operates through NCLX.

Unexpectedly, the downstream effects of Ca^2+^ release are not primarily directed at mitochondrial fission. Although cytosolic Ca^2+^ has been shown to activate Drp1 through CaMK-dependent phosphorylation (Han *et al*., 2008; Xu *et al*., 2016) or calcineurin-mediated dephosphorylation (Cereghetti *et al*., 2008; Cribbs & Strack, 2007), inhibiting NCLX caused only minor changes in stress-induced fission. In contrast, blocking AMPK significantly reduced fission, aligning with ATP depletion and AMPK-dependent phosphorylation of MFF (Toyama *et al*., 2016). Therefore, mitochondrial ROS-induced fission appears mostly independent of mitochondrial Ca^2+^ efflux and instead may result from ATP depletion.

ROS-induced Ca^2+^ release may affect stages of mitophagy other than fission. We identify the Ca^2+^-regulated E3 ubiquitin ligase NEDD4-1 as a key effector necessary for progressing ROS-induced mitophagy. NEDD4-1 has a Ca^2+^-binding C2 domain that binds to PI(3)P-enriched membranes and an LC3-interacting region that helps it engage with the autophagic machinery (Sun *et al*, 2017). NEDD4-1 polyubiquitinates SQSTM1/p62 (Lin *et al*, 2017), boosting its receptor activity and stabilizing protein complexes with autophagic cargo (Kumar *et al*, 2022; Peng *et al*, 2017; Xiao *et al*, 2025). NEDD4-1 might work together with other E3 ubiquitin ligases that initiate mitophagy by ubiquitinating mitochondrial outer membrane proteins (Wang *et al*, 2026). Evidence supporting a pro-mitophagic role of Ca^2+^ release comes from OPA1 heterozygous mice, which show chronic mitochondrial Ca^2+^ release, excessive mitophagy, and cell death (Zaninello *et al*, 2022). This may be due to impaired NCLX retention within cristae and increased Mfn2-NCLX interactions. Although NCLX-dependent Ca^2+^ signaling has been linked to actin polymerization and interference with Parkin-dependent mitophagy after depolarization (Chakrabarti et al., 2022; Fung et al., 2025), our ROS-inducing conditions keep the mitochondrial membrane potential intact, pointing to a pathway distinct from depolarization-driven mitophagy.

In summary, our results support a model in which mitochondrial ROS trigger Ca^2+^ release through the coordinated regulation of Mfn2, SLC25A46, and NCLX. The resulting Ca^2+^ signal does not mainly cause fission but instead promotes mitophagy, establishing a distinct stress-response pathway originating from mitochondria. The novel role of Mfn2 as a regulator of mitochondrial Ca^2+^ release may have implications for diseases caused by Mfn2 mutations, such as Charcot-Marie-Tooth disease type 2A and cardiomyopathy, in which mitophagy defects have also been observed (Franco *et al*, 2023). This indicates a shared mechanism linking Mfn2, Ca^2+^ balance, and disease development. These defects may occur simultaneously or worsen existing issues with mitochondrial fusion, transport (Stuppia, 2015 #4401; Zhou, 2019 #5159), and ER-mitochondria tethering (Bernard-Marissal, 2019 #4946) caused by mutations in Mfn2. By showing how mitochondrial ROS is converted into a cytosolic Ca^2+^ signal that guides selective mitochondrial turnover, this work provides a mechanistic link between oxidative stress, organelle communication, and mitochondrial quality control.

## Materials and methods

### Antibodies and chemicals

Rabbit polyclonal anti-Mfn1, NCLX, SLC25A46, and Tomm20 antibodies were from ProteinTech. Rabbit polyclonal anti-Mfn2 and Oma1 antibodies were from Cell Signaling. Rabbit polyclonal anti-Vinculin and Myc antibodies were from Sigma. Mouse monoclonal anti-Tubulin and Actin antibodies were from Sigma. Mouse monoclonal anti-CNX, Opa1 and Drp1 antibodies were from BD Pharmingen. Mouse monoclonal anti-Hsp60 antibodies were from Abcam. Mouse monoclonal anti-FLAG antibodies were from Proteintech. Mouse monoclonal anti-HA antibodies were from Millipore. Chicken polyclonal ant-Hsp60 antibodies were from EnCor. Oligomycin, Antimycin, CGP37157, CCCP, TMRM, DSP, SPDP and Dodecyl-maltoside were from Sigma. PXA was from Adipogene. Rhod-2AM was from Biotium. FLAG Magnetic Beads were from Pierce.

### DNA and RNA

The Mfn2-16xmyc, pCMV R-Cepia3mt and px459 plasmids were from Addgene (#23213, #140464 and #62988, respectively). The pPB EF1A>hNCLX-3xHA and pPB EF1A>hMfn2-3xFlag and pBase plasmids were from Vectorbuilder. The pCDNA3 hMfn2-FLAG plasmid was a kind gift from Gerald Dorn (Washington University). Target sites for gene deletions were identified using the Boutros Lab Website (http://www.e-crisp.org/E-CRISP/designcrispr.html) or as recommended by the Vectorbuilder website. The gRNAs were cloned in the px459 plasmid or in a custom vector from Vectorbuilder. The gRNAs used in this study are listed in Table 1. NCLX expression was knocked down in MEFs with shRNA as described (Palty *et al*., 2010) and in HeLa cells with mixture of 2 predesigned Dicer-Substrate siRNAs from IDT (cat nr. hs.Ri.SLC8B1.13.2 and hs.Ri.SLC8B1.13.3. Site-directed mutagenesis used the DpnI-method for double-strand plasmids(Braman *et al*, 1996; Kolitsida *et al*, 2019).

### Cell culture, transfections and gene knockouts

HeLa cells were from James Wohlschlegel (Dept. of Biological Chemistry, UCLA) and MEFs were from David Chan (Dept of Biology, CalTech). Cells were grown in glycolytic medium (DMEM with 10% FBS with 1% Pen/Strep and 4.5 g/l glucose) or respiring medium (DMEM without glucose and supplemented with 4.5g/L galactose, 10% FBS, 1% PS, 5mM sodium pyruvate and 2mM L-glutamine) as indicated. All cell lines were periodically checked for Mycoplasm. Transient transfections were done with jetPRIME following manufacturer’s instructions (Polyplus). For siRNA, cells were grown in 6cm dishes, transfected with 50nM oligonucleotides using RNAimax (Invitrogen) and analyzed 72h later. For gene knockouts, 1.2μg/well was transfected into 6 well plates cells, followed by selection at increasing concentrations or puromycin from 1µg to 5 µg/ml for 1-2 days. Surviving colonies were isolated and analyzed with Western blots. A stable cell line that expresses human Mfn2 with 3XFLAG under the EF1alpha promoter was generated with a PiggyBac construct from Vectorbuilder. This construct was transfected into Drp1/Mfn2 DKO MEFs along with the PBase vector from Vectorbuilder at equal concentrations (2µg/per well in a 6 well plate). At 24h post transfection, cells were subjected to puromycin selection at increasing concentrations from 1µg to 5 µg/ml for 1-2 days.

After selection, surviving colonies were isolated with cloning rings, expanded and analyzed for Mfn2 expression levels by Western blotting.

*<u>For lentiviral transduction</u>*, cells were seeded in 6-well plates at a density of 1.5–2 × 10⁵ cells per well to achieve 50–70% confluency at the time of infection. On the day of transduction, the culture medium was replaced with fresh complete medium containing polybrene (5 µg/mL final concentration) to enhance viral entry. A concentrated lentiviral stock for mitoQC (Welgen Inc) was diluted 1:1000 in complete medium and added dropwise to the cells. Cells were incubated with the virus-containing medium for 48 h at 37 °C. After this period, the viral supernatant was removed and fresh complete medium without polybrene was added. Transduction efficiency was assessed 72 h post-infection by fluorescence microscopy for GFP/RFP expression.

### Immunoblotting and immunofluorescence

Total cell lysates for Western blotting were prepared in RIPA buffer. Samples were subjected to SDS-PAGE, transferred to PVDF membranes, blocked with 5% non-fat milk, and incubated overnight at 4 °C with primary antibodies. Membranes were then washed with TBS- T and incubated with secondary antibodies. Chemiluminescent bands were detected with a BioRad scanner. For immunofluorescence images, cells were grown on 12mm coverslips, fixed for 15min with 4% paraformaldehyde in PBS, and permeabilized for 15min with 0.25% Triton X-100 in PBS, blocked for 1h with BSA in PBS-T and incubated with primary antibodies. Secondary antibodies were Alexa Fluor 488-, 594- or 647-conjugated goat anti-mouse or rabbit IgG (Invitrogen).

<u>Proximity ligation assays</u> were conducted with Duolink as recommended by the manufacturer (Sigma-Aldrich). Cells were seeded on coverslips and at 60% confluency they were transfected with Mfn2-myc and NCLX-HA overnight. The cells were then treated for 5-10 min with 10µM PXA or for 30min 10µM oligomycin or DMSO as indicated, after which they were processed for PLA as described (Alam, 2018), but with the following modification: After the fixation and washing, cells were permeabilized with 0.3% Triton-X100 for 10 minutes at room temperature and then washed twice for 5 min with PBS. Along with the primary antibodies for PLA, we also added chicken anti-HSP60 antibody (EnCor Biotechnology Inc) or TOM20 (Proteinteck) and anti-chicken or anti rabbit Alexa Fluor 488 as secondary antibody, along with the primary and secondary antibodies for PLA.

*<u>Microscopy</u>* was performed with a Marianas spinning disc confocal from Intelligent Imaging, which uses an Axiovert microscope (Carl Zeiss Microscopy) with 40x/1.4 and 100x/1.4 oil objectives, a CSU22 spinning disk (Yokogawa), an Evolve 512 EMCCD camera (Photometrics), and a temperature unit (Okolab). For live cell imaging, cells were grown in glass bottom dishes (MatTek) and imaged in situ at 37°C. Fiji software was used to determine Manders’ coefficients and generate scatter plots. Aspect ratios of mitochondria were determined with Tomm20 fluorescence images. First, cells in the images were demarcated using FIJI ImageJ (https://fiji.sc/). Mitochondria were then demarcated using CellProfiler (https://cellprofiler.org/), and background noise was suppressed with a median filter. Mitochondria were segmented with a global, three-class, Otsu-thresholding method, minimizing the weighted-variance to shape. The aspect ratio of each object was then determined as the quotient of Major Axis Length over Minor Axis Length. Data represent the average aspect ratio for 25 cells in 3 independent experiments using WT and Oma1 KO HeLa cells.

*Mitochondrial Ca^2+^ levels* were determined in HeLa cells with transiently transfected pCMV R-Cepia3mt (Addgene #140464). Transfected cells were first perfused for 30 min with Krebs-Ringer solution to record a baseline signal. Where indicated 10 μM CGP37157 was included with the perfusion. Mitochondrial Ca^2+^ release was then triggered by adding 10 μM PXA, 10 μM Oligomycin or 50 μM mitoPQ. Using 3I software, fluorescence intensities of individual cells was determined once per second over a period of 10 min after the addition of PXA or Oligomycin. Mitochondrial calcium levels in MEFs were measured using a Tecan Spark 10M multimode plate reader equipped with an injector, as previously described (Martinez *et al*, 2017). In brief, MEF DRP1 KO or double MFN2/DRPI KO cells were plated on Corning 96 Flat black, clear bottom wells and loaded with 1 μM Rhod2-AM for 30 min at 37 °C using a modified Krebs–Ringer’s solution containing (126 mM NaCl, 5.4 mM KCl, 0.8 mM MgCl_2_, 20 mM HEPES, 1.8 mM CaCl_2_, 15 mM glucose, with pH adjusted to 7.4 with NaOH and supplemented with 0.1% BSA). After dye loading, cells were washed three times with fresh dye-free Krebs–Ringer’s solution, followed by additional incubation of 30 min to allow for the de-esterification of the residual dye. Kinetic live-cell fluorescent imaging was performed to monitor Ca^2+^ transients. Rhod2-AM was excited at 552 nm wavelength light and imaged with a 570 nm. After establishing a baseline, cells were triggered with PXA at a final concentration of 10 μM. Kinetic measurements were taken at ∼ 5 s intervals. Traces of Ca^2+^ responses were analyzed and plotted using KaleidaGraph. The rate of ion transport was calculated from each graph (summarizing an individual experiment) by a linear fit of the change in the fluorescence over time (Δ*F*/d*t*), as previously described (Assali *et al*, 2020; Taha *et al*, 2024).

*Cytosolic Ca^2+^ levels* in HeLa cells were determined after loading with Calbyte-520aAM for 30min at RT using a Krebs–Ringer’s solution containing (126 mM NaCl, 5.4 mM KCl, 0.8 mM MgCl_2_, 20 mM HEPES, 1.8 mM CaCl_2_, adjusted to pH 7.3 with NaOH). After dye loading cells were washed three times with PBS, followed by additional incubation of 30 min to allow for the de-esterification of the residual dye. Calbryte 520-AM was excited at 488 nm wavelength light and imaged with a 520 nm. These transfected cells were first perfused for 30 min with Krebs-Ringer’s solution included the treatment 10 μM CGP37157. Cytosolic Ca^2+^ release was then triggered by adding 10 μM Oligomycin or 50μM mitoPQ. Using 3I software, fluorescence intensities of individual cells was determined once per second over a period of 10 min after the addition of oligomycin or mitoPQ.

*<u>Mitochondrial membrane potential</u>* was detected with TMRM (Thermo-Fisher Scientific). Cells plated in 35 mm glass bottom dishes (MatTek) were rinsed with 10 mM HEPES buffered HBSS, pH 7.3 with 15 mM glucose and subsequently incubated with 25 nM TMRM for 30 min at 37°C, followed by 3 washes with PBS and destaining for 30 min at 37°C in 1.8mM CaCl_2_, 120mM NaCl, 5.4mM KCl, 0.8mM MgCl_2_, 20mM HEPES, 15mM glucose, adjusted to pH 7.3 with NaOH. Fluorescence images were acquired for a field of cells with 10x objective. Where indicated, cells were treated for 30 min at 37°C with 10 μM CGP37157 before adding 10 μM PXA. As a reference for uncoupling, cells were treated with 10 μM CCCP.

### Cellular Thermal Shift Assay (CETSA)

MEFs cultured to 1.0 × 10^7^ cells in a 15 cm dish were washed with PBS and harvested in 2 ml lysis buffer (1.8mM CaCl2, 120mM NaCl, 5.4mM KCl, 0.8mM MgCl2, 20mM HEPES, 15mM glucose, 0.5% Dodecyl-Maltoside, adjusted to pH 7.3 with NaOH). Equal volumes of cell lysates were transferred to 1.5 ml Eppendorf tubes for incubation with 1 μM PXA or DMSO for 4 min at RT. After this incubation, 100 μl aliquots of cell lysates were transferred to PCR tubes and heated on a gradient PCR machine for 3 min at different temperatures as indicated and then cooled to 25°C for 2 min, followed by centrifugation for 30 min at 20.000g in 4 °C. Supernatants were then transferred to new tubes with sample buffer. For each temperature, 10 μl (approximately 30 μg protein) was analyzed by SDS-PAGE and western blotting. Band intensities were quantified with densitometry and ImageJ.

### Co-immunoprecipitation

MEFs stably expressing Mfn2-FLAG were treated for 5 minutes at 37 °C with DMSO, 10µM PXA, 10µM CCCP, or 10µM CGP37157 as indicated. These cells were then washed with ice cold PBS with 0.9 mM CaCl_2_ and 0.5 mM MgCl_2_ and crosslinked for 2h on ice with 1 mM DSP or 1 mM BMH. The crosslinkers were then quenched by incubating for 15 minutes on ice with 20 mM Tris-Cl (pH7.4) followed by a wash with ice cold PBS with 0.9 mM CaCl_2_ and 0.5 mM MgCl_2_ and resuspending in RIPA buffer. Co-immunoprecipitations were conducted with 1 mg protein (determined with BCA assay). 50 μl of magnetic FLAG beads (Pierce/Sigma) per sample were washed twice with 1X TBST and then incubated with protein samples for 45min at 4 °C on a rotor. Beads were then collected using a magnetic stand and the supernatant was discarded. The beads were washed twice with 1X TBST and then once with ddH2O. Finally, the beads were resuspended in 50 µl of 2X sample buffer, heated for 5 min at 95 °C, and analyzed by western blot.

### Statistics & reproducibility

All statistical analyses were performed on Graphpad Prism Version 8.4.1. All column graphs were generated on Graphpad Prism Version 8.4.1. Data points from all biological replicates were tested for multiple comparison using Turkey test and ANOVA test. The bars indicate standard deviations. Each data point on a column graph represents one biological replicate. All quantitative analysis was performed on at least three biological replicates per condition. Data are represented as mean, and error bars indicate the standard deviation of the mean.

## Acknowledgements

CMK was supported by NIH grants R01GM61721, R01GM073981 and R01DK101780. AMvdB was supported by NIH grants U01GM109764 and R01NS120690.

---

**Fig. S1.**
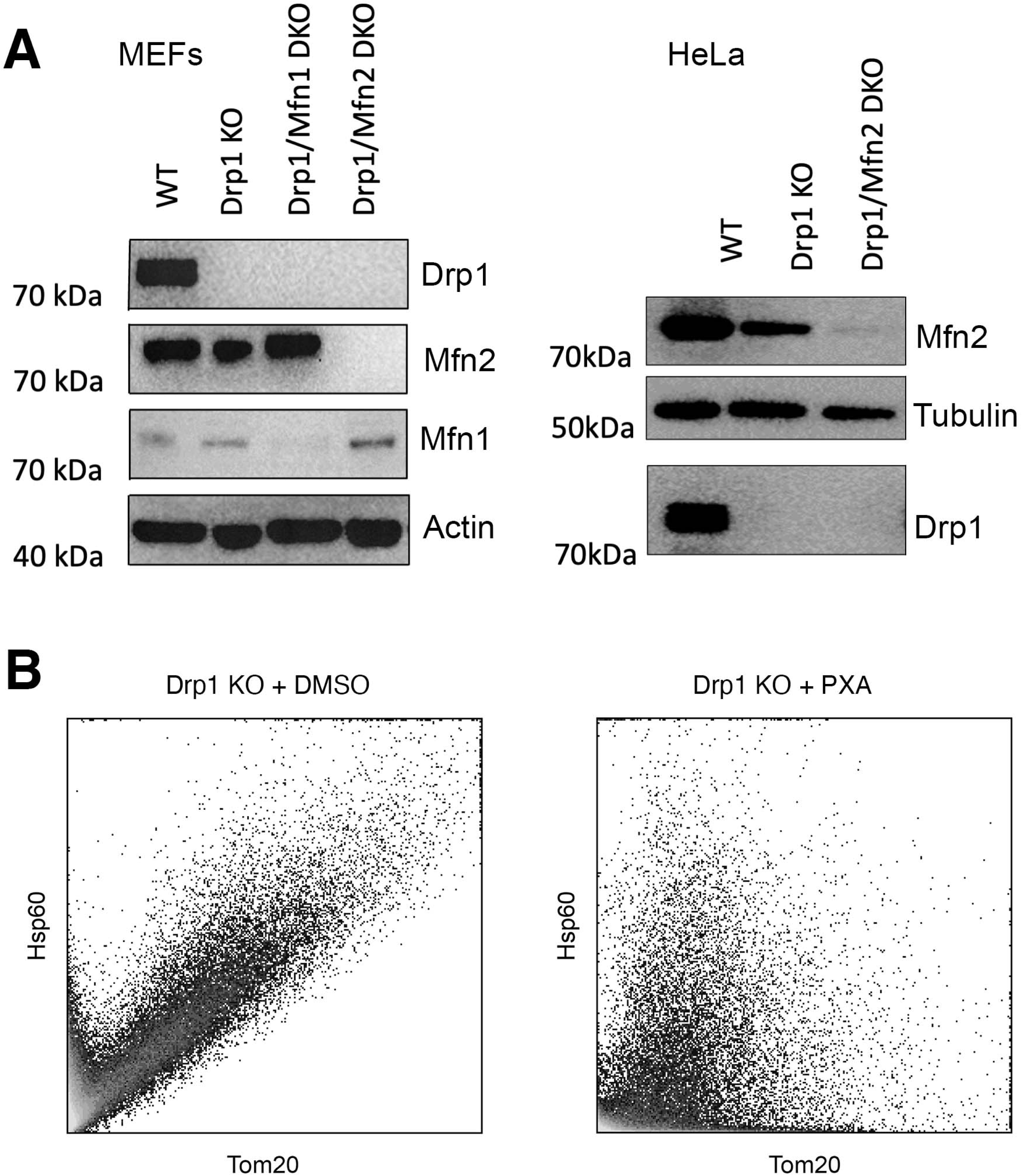
Western blots of knockout cells and scatterplots for immunofluorescence analysis. (**A**) Western blot confirmation of knockouts used for IF experiments. Actin and tubulin are loading controls. (**B**) Scatter plots of Hsp60 and Tom20 immunofluorescence with Drp1 KO cells treated with or without 10 μM PXA to illustrate colocalization as an indicator of mitochondrial matrix constrictions. Matrix and outer membrane markers largely colocalize in cells treated with DMSO, as evidenced by a linear relation in the scatterplot, whereas they separate when treated with PXA, resulting in a random distribution in the scatterplot.

**Fig. S2.**
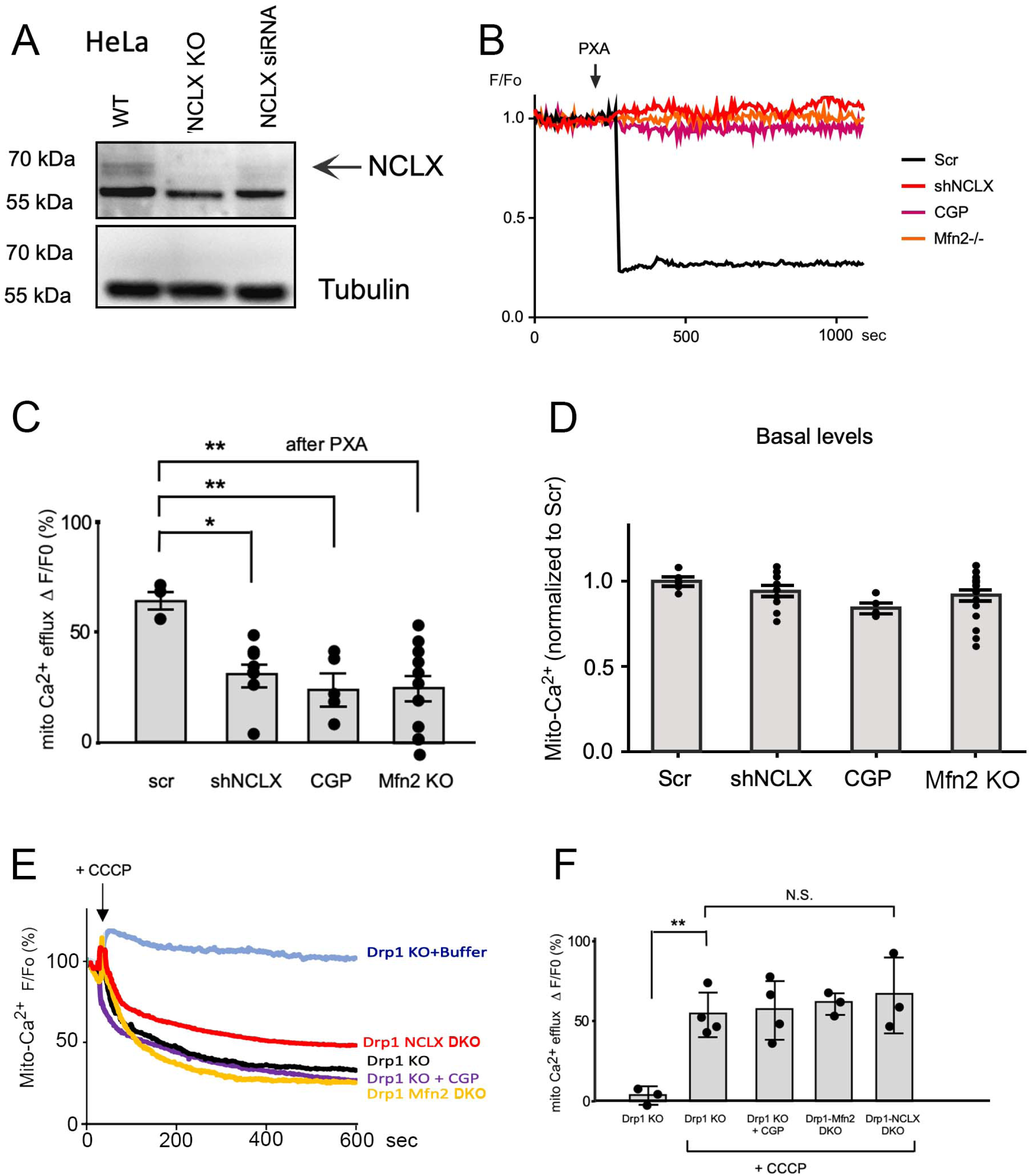
Western blots showing NCLX knockout in HeLa cells, PXA-induced Ca^2+^ release detected with Rhod-2AM, and plots of mitochondrial membrane potential affected by Ca^2+^ release. (**A**) Validation of NCLX KO in HeLa cells with a western blot of extracts from NCLX KO cells, flanked by extracts from WT and NCLX siRNA cells. The NCLX antibody cross-reacts with another protein at around 57 kDa, but a band corresponding to the predicted MW of NCLX (64 kDa, shown with an arrow) is present in WT cells, absent from NCLX KO cells, and greatly diminished in NCLX siRNA cells. (**B**) Time course of soluble mitochondrial Ca^2+^ levels in MEFs detected with Rhod-2 AM, upon treatment with 10 μM PXA. Drp1 KO MEFs were transduced with scrambled siRNA (SCR), NCLX siRNA, or pretreated for 30 min with 10 μM CGP37157. Mfn2-Drp1 DKO MEFs were also tested. (**C, D**) Quantification of mitochondrial Ca^2+^ levels detected with Rhod-2 AM as in panel B, showing induced and basal levels. (**E**) Tracing of mitochondrial Ca^2+^ levels in HeLa cells induced by 10 μM CCCP and detected with matrix-3mt-CEPIA as in Fig. 2A, showing that CCCP induces Ca^2+^ release independent of NCLX function. (**F**) Quantification of mitochondrial Ca^2+^ levels after CCCP treatment. Throughout, averages are shown with SD and significance was determined with one-way ANOVA with Tukey’s HSD post hoc test.

**Fig. S3.**
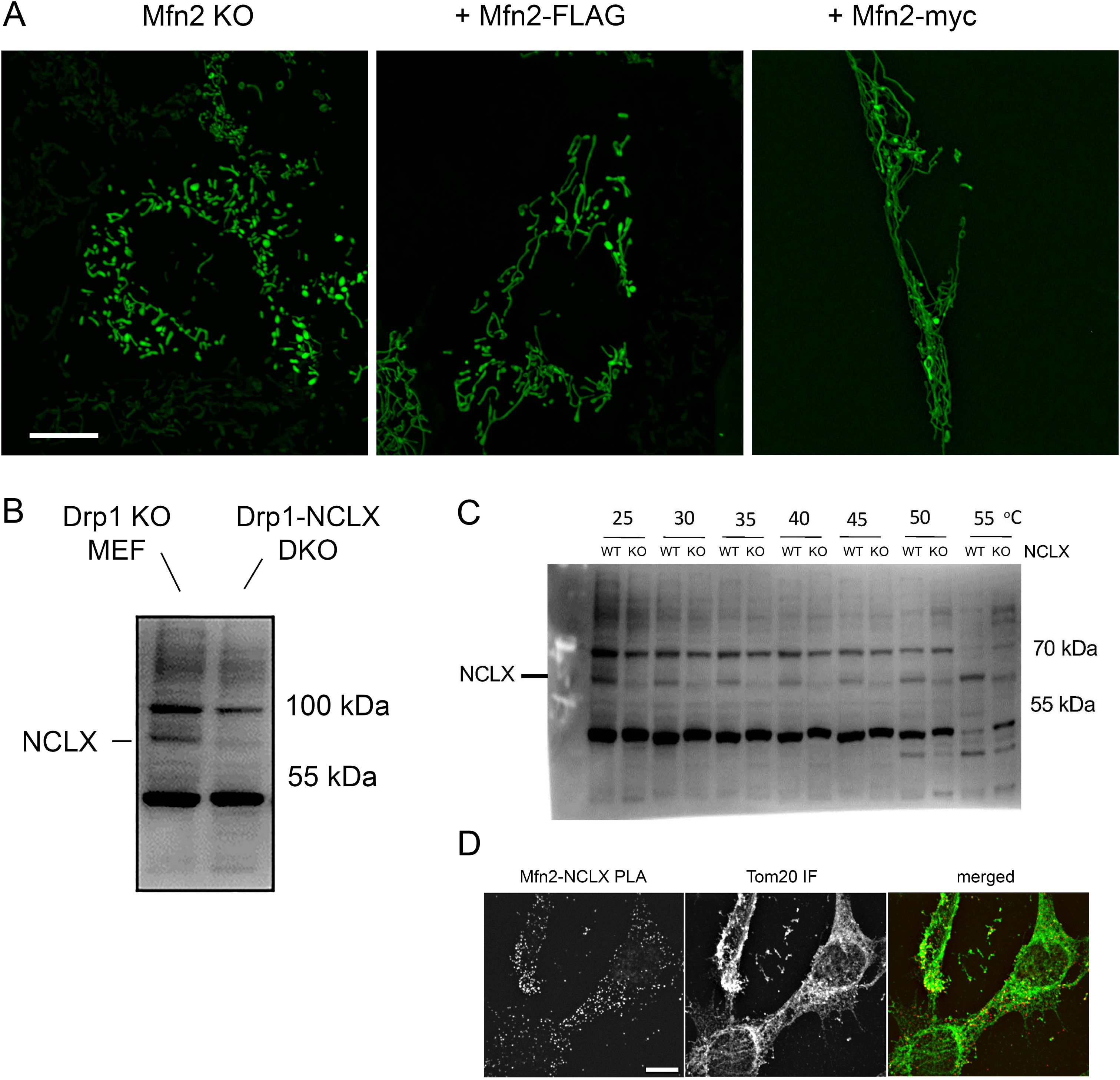
Verification that Mfn2-flag and Mfn2-myc tagged proteins are functional. (**A**) Fluorescence images showing that stable transfection with tagged Mfn2 constructs (Mfn2-FLAG or Mfn2-myc) restores filamentous mitochondrial morphology in Mfn2 KO MEFs. Mitochondria were detected with mito-GFP. Scale bar is 10 μm. (**B**) Identification of NCLX band in MEF extracts under conditions for CETSA experiments. This was necessary because the cross-reactivity observed with the available NCLX antibody changes with different solubilization conditions (see westerns with Laemmli sample buffer in HeLa cells (Fig. S2) versus dodecyl maltoside for the MEFs used for CETSA (Fig. 3). In both cases, NCLX was identified by the absence of a 64 kDa band in KO or siRNA cells, which is the predicted MW of NCLX. (**C**) Example of CETSA blot without drugs, but with extracts from WT and NCLX KO MEFs to show the stability over a range of temperatures and positive identification of the NCLX band by comparing with extracts from KO cells. (**D**) PLA of Mfn2-myc and NCLX-HA (red spots) in glycolytic Drp1 KO MEFs treated with DMSO. In this case mitochondria were detected by immunofluorescence with an antibody against a mitochondrial outer membrane protein (rabbit Tom20 antibody, green), showing that the spots are outer membrane tubules. Size marker is 10 μm.

**Fig. S4.**
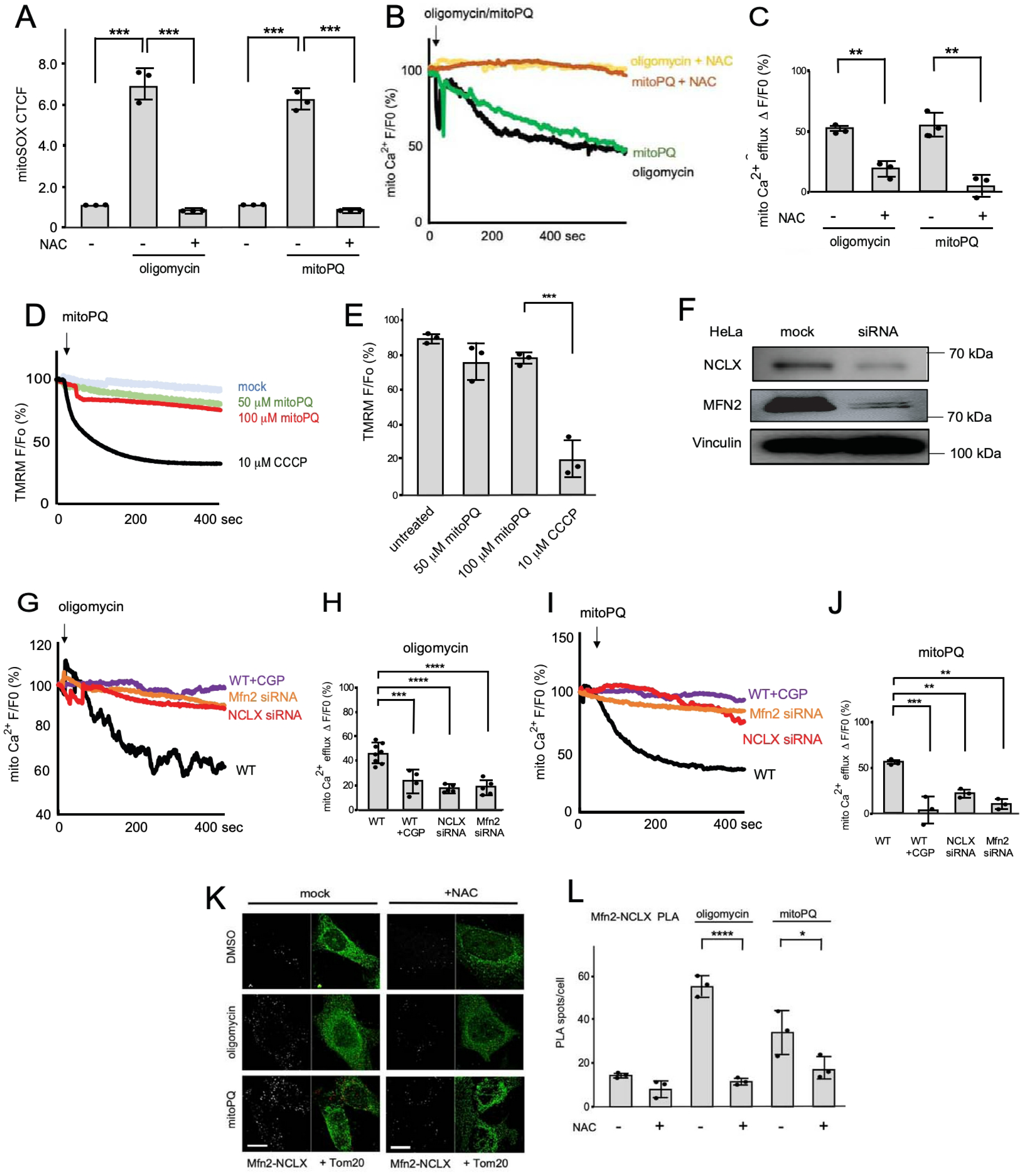
Mitochondrial ROS, Ca^2+^ release and membrane potential in different mutant HeLa cells. (**A**) Mitochondrial ROS detected with mitoSOX and expressed as corrected total cell fluorescence (CTCF) values. ROS was induced by oligomycin or mitoPQ, and as a control, quenched with NAC. ROS was detected by incubating WT HeLa cells for 30 min with 5 μM MitoSOX, with or without 30 min pre-treatment with 2 mM NAC. After this incubation, cells were washed with PBS and destained for 30 min during which cells were treated with 10 μM Oligomycin or 50 μM mitoPQ. CTCF was quantified per cell (25 cells per condition) across 3 independent experiments. (**B**) Tracings of mitochondrial matrix Ca^2+^ in respiring HeLa cells, detected as in Fig. 2A, but induced with 10 μM oligomycin or 50 μM mitoPQ, with or without pretreatment for 30 min with 2mM NAC. (**C**) Changes in mitochondrial Ca^2+^ levels reflected by the relative fluorescence (F/Fo) at 600 sec. (**D**) Effects of 50 μM or 100 μM mitoPQ on mitochondrial membrane potential (ΔΨ) determined with WT HeLa cells loaded with 25 nM TMRM. (**E**) Averages of F/Fo for TMRM determined at 400 sec after treatment with mitoPQ. (**F**) Validation of NCLX and Mfn2 siRNA knockdowns in HeLa cells with western blots. Vinculin is a loading control. (**G**) Tracings of mitochondrial matrix Ca^2+^ in respiring HeLa cells, detected as in Fig. 2A, but induced with 10 μM oligomycin. Where indicated, cells were pretreated for 30 min with 10 μM CGP37157 or transfected with siRNA. (**H**) Changes in mitochondrial Ca^2+^ levels reflected by the relative fluorescence (F/Fo) at 600 sec. (**I, J**) Tracings and histograms of mitochondrial matrix Ca^2+^, as in panels A and B but treated with 50 μM mitoPQ. Throughout, averages are shown with SD and significance was determined with one-way ANOVA with Tukey’s HSD post hoc test. (**K**) PLA of Mfn2-myc and NCLX-HA (red spots) in respiring Drp1 KO MEFs pre-treated for 30 min with 2mM NAC and then incubated for 30 min with DMSO, 10 μM oligomycin, or 50 μM mitoPQ. Mitochondria were detected with chicken Tom20 antibody (green). The size marker is 10 μm. (**L**) Number of PLA spots for Mfn2-myc and NCLX-HA per cell, averaged across 25 cells for each condition in three rounds of experiments.

**Fig. S5.**
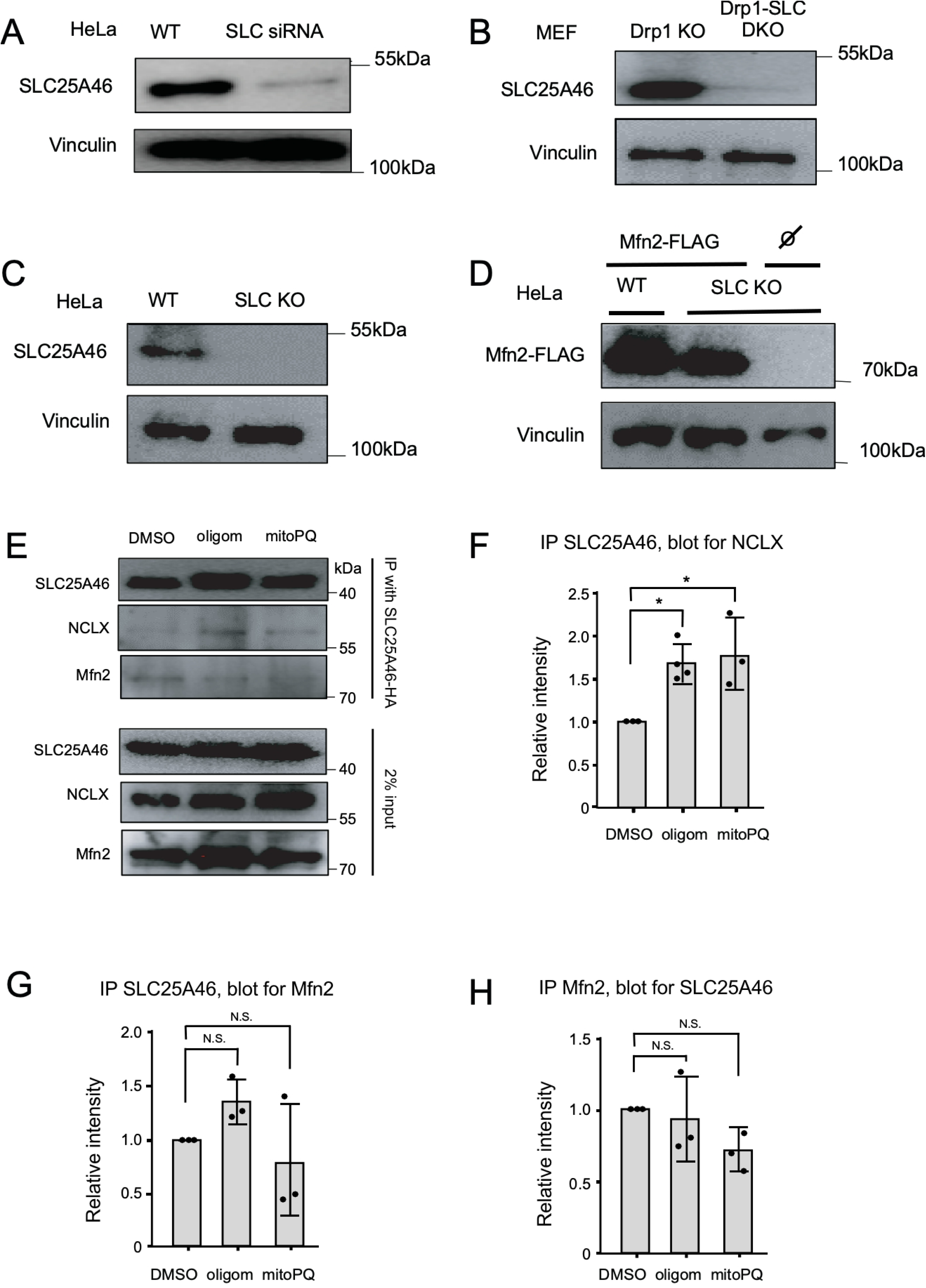
Western blots to confirm SLC25A46 siRNA and knockout in HeLa cells and MEFs. (**A**) Validation of SLC25A46 siRNA knockdown in HeLa cells with western blots of extracts from WT and siRNA-treated cells. Vinculin is a loading control. (**B**) Validation of SLC25A46 KO in Drp1 KO MEFs. (**C**) Validation of SLC25A46 KO in HeLa cells. (**D**) Comparison of Mfn2-FLAG expression levels in WT and SLC25A46 KO HeLa cells. (**E**) Co-immunoprecipitation (coIP) of endogenous NCLX and Mfn2 with SLC25A46-HA using HA antibody beads. Drp1 KO MEFs grown under respiring conditions were treated for 30 min with DSMO, 10 μM oligomycin, or 50 μM mitoPQ, followed by incubation with BMH cross-linker, coIP and western blot analysis. (**F**) Densitometry of the NCLX coIP with SLC25A46, normalized to the levels of SLC25A46 for each condition. (**G**) Densitometry of Mfn2 coIP with SLC25A46, normalized to the levels of SLC25A46 for each condition. (**H**) Densitometry of SLC25A46 coIP with Mfn2, normalized to the levels of Mfn2 for each condition. Throughout, averages are shown with SD and significance was determined with one-way ANOVA with Tukey’s HSD post hoc test.

**Fig. S6.**
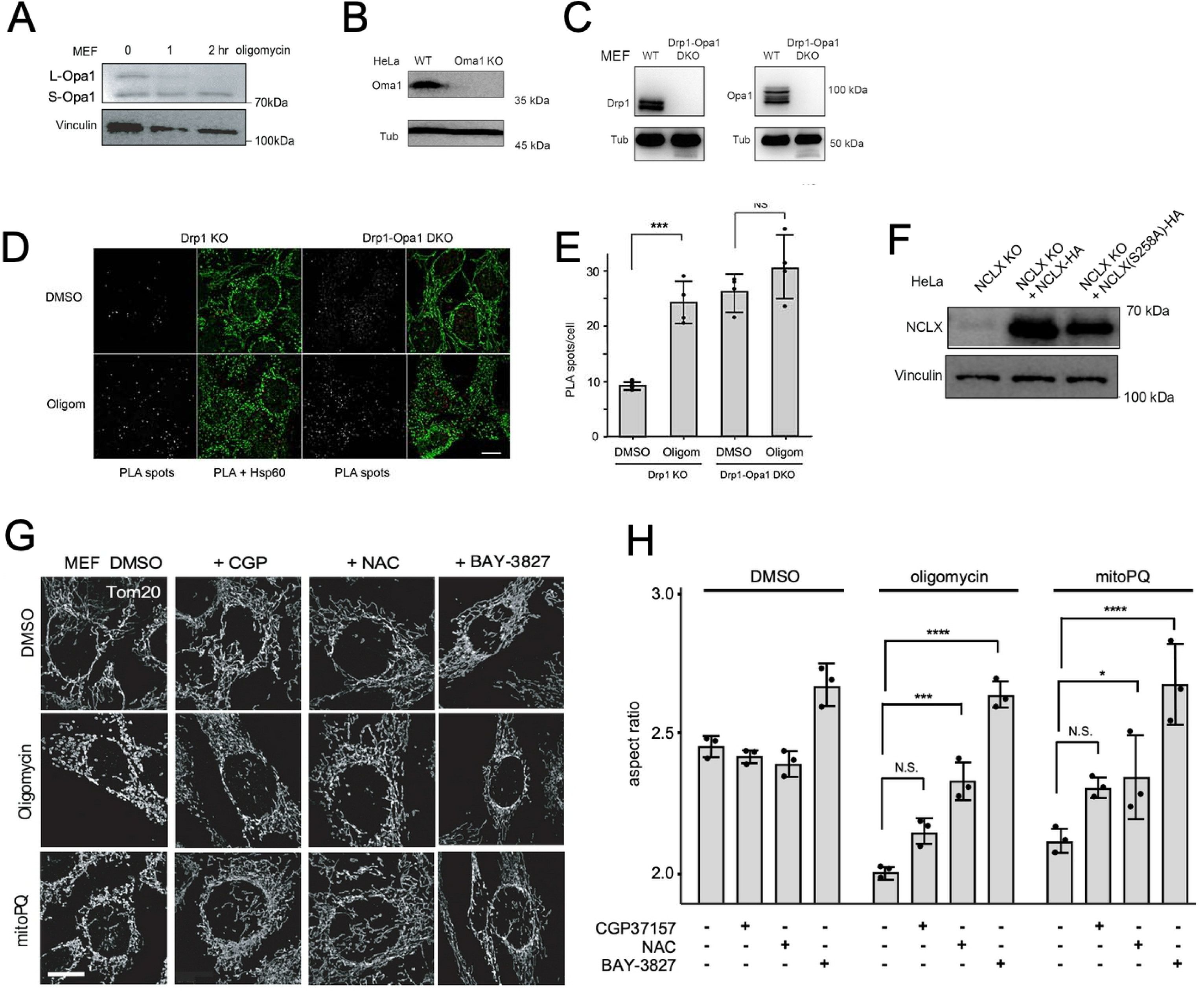
Oma1 cleavage of Opa1 controls Mfn2-NCLX association and Ca^2+^ release. (**A**) Western blot analysis of WT MEF cells treated with 10 µM Oligomycin for 0, 1, or 2 hours at 37 °C, probed with an Opa1 antibody. Tubulin was a loading control. (**B**) Western blot of wild type (WT) and Oma1 KO HeLa cells. Tubulin was a loading control. (**C**) Western blot of WT and Drp1-Opa1 DKO MEFs. (**D**) PLA of Mfn2-myc and NCLX-HA (red spots) in Drp1 KO and Drp1/Opa1 DKO MEFs grown as in panel D under respiring conditions and treated for 30 min with DMSO or 10 μM oligomycin. Mitochondria were detected with a chicken anti-Hsp60 antibody (green). Scale bar is 10 μm. (**E**) Numbers of PLA spots per cell in 25 cells treated with DMSO or 10 μM oligomycin from 3 independent experiments with SD and results of a one-way ANOVA. (H) Western blot showing stable expression of NCLX(WT)-HA and NCLX(S258A)-HA in HeLa NCLX KO cells. (**F**) Western blot of NCLX KO HeLa cells with stable expression of WT NCLX-HA or NCLX^S258A^-HA. Vinculin is a loading control. (**G**) WT MEFs were treated with DMSO or 10 μM CGP37157 or 5 mM NAC for 30 min or 10 μM BAY-3827 for 2h at 37 °C followed by 10 μM Oligomycin or 50 μM mitoPQ for 30 min at 37 °C and labeled with Tom20 antibody for immunofluorescence. The size marker is 10 μm. (**H**) Aspect ratios of mitochondrial outer membrane (Tom20) under the conditions for panel a. Averages are shown for 3 independent experiments with SD and the results of one-Way ANOVA. Aspect ratios were determined for each condition with mitochondria in 25 cells per experiment.

**Fig. S7.**
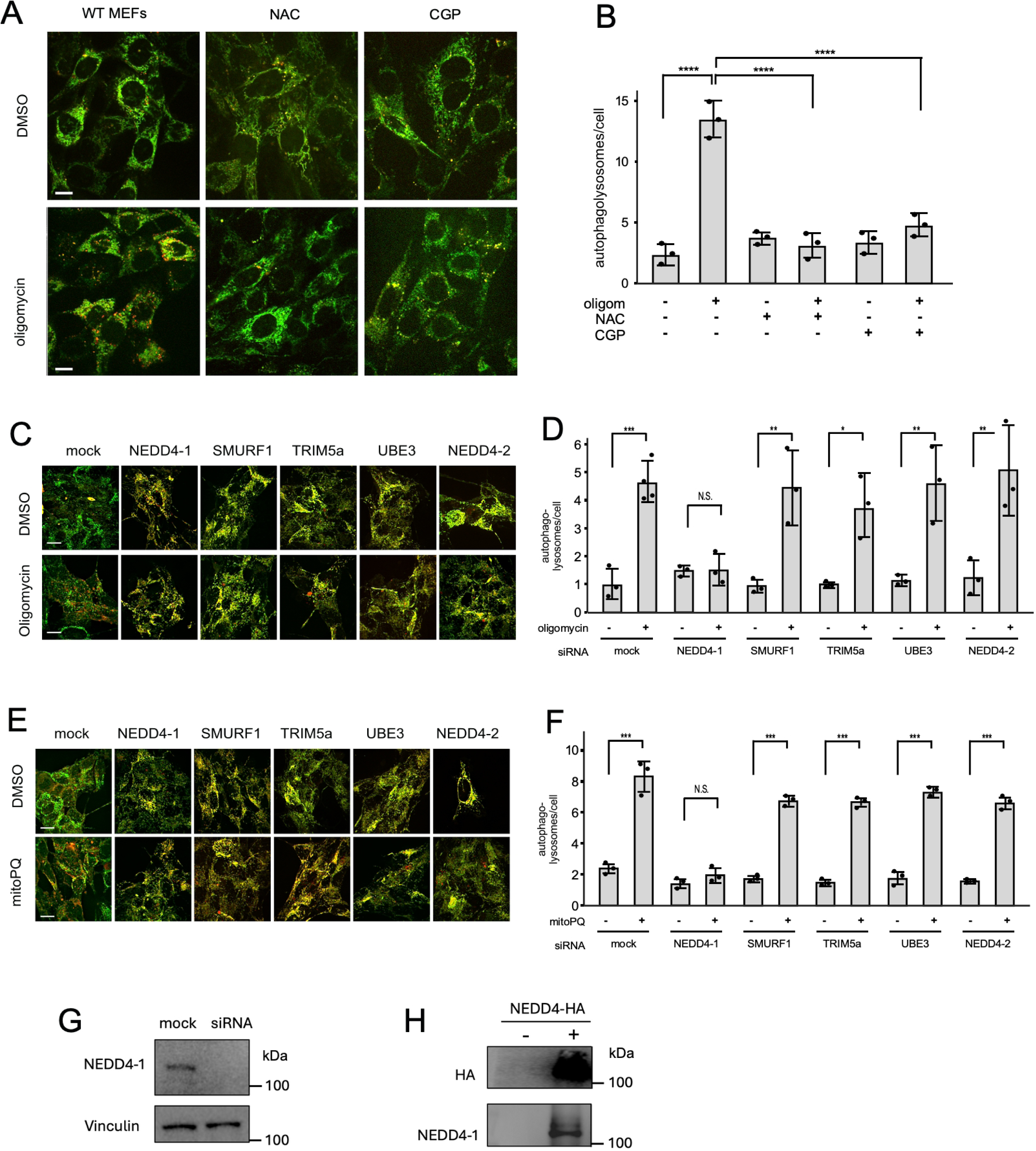
Effects of ROS-induced mitochondrial Ca^2+^ release on mitophagy and the effects of Ca^2+^ dependent E3 ubiquitin ligases on this process. (**A**) Representative live images of WT MEFs that stably express mitoQC (Fis1-mCherry-GFP) introduced with a lentivirus. Cells were pre-incubated with 10 μM CGP37157 or 5 mM NAC for 30 min at 37 °C and treated for 10 μM oligomycin for 30 min at 37 °C. Scale bar is 10 μm. (**B**) Quantification of red puncta per cell from 3 independent experiments with SD and results of one-Way ANOVA with Tukey’s HSD post hoc test. (**C**) Live cell images of SH-SY5Y cells that express mitoQC, transfected with siRNA for genes as indicated, followed after 2 days by inducing mitophagy with 10 μM oligomycin for 30 min at 37 °C. Scale bar is 10 μm. (**D**) Quantification of red puncta per cell from 3 independent experiments with SD and results of one-Way ANOVA with Tukey’s HSD post hoc test. (**E, F**) SH-SY5Y cells that express mitoQC were transfected with siRNA for the indicated genes as in panels C and D, but these were treated for for 30 min with 50 μM mitoPQ. (**G**) Western blot showing knockdown of NEDD4-1 with siRNA. (**H**) Western blot showing NEDD4-1 expression in cells transfected with an HA-tagged NEDD4-1 construct.

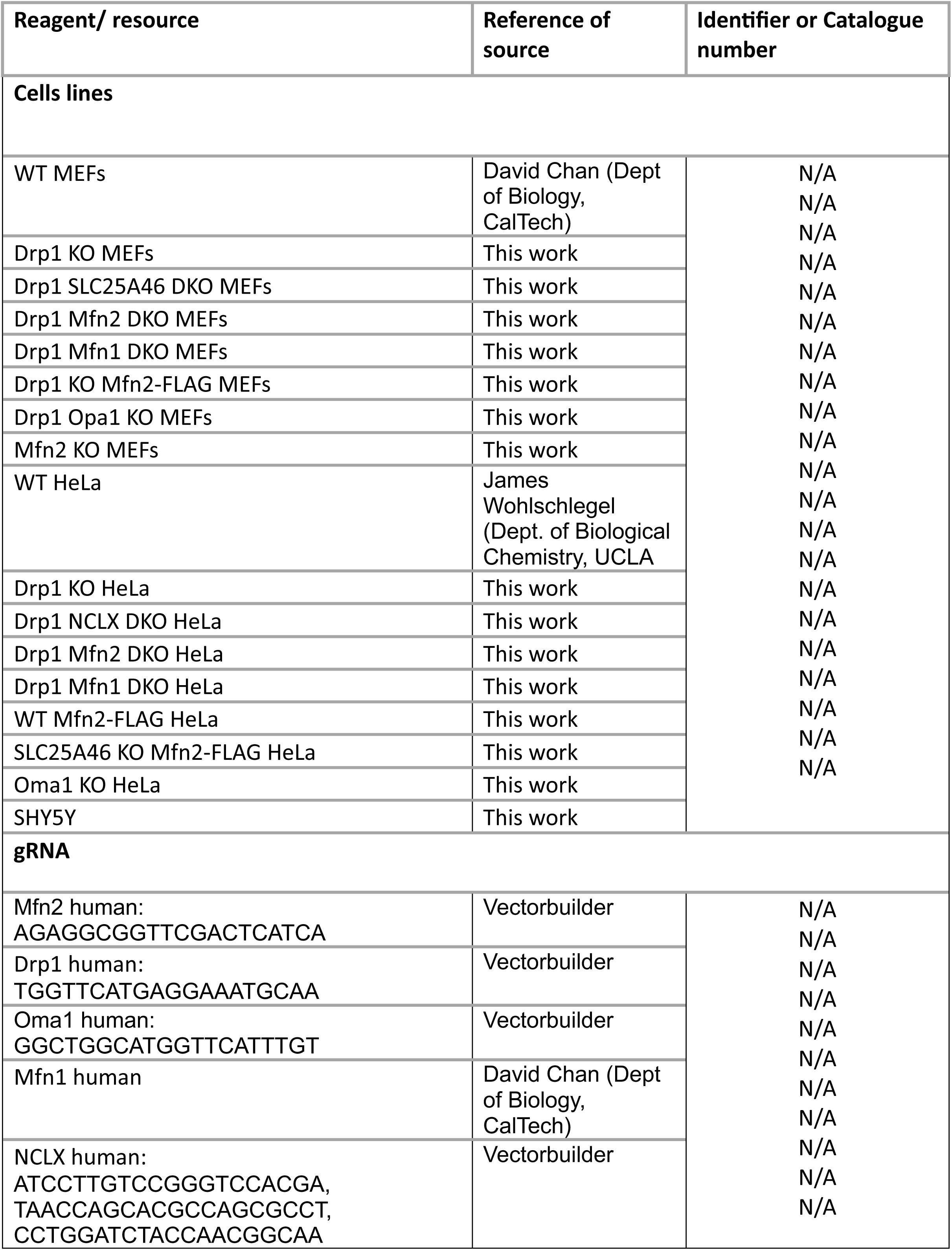
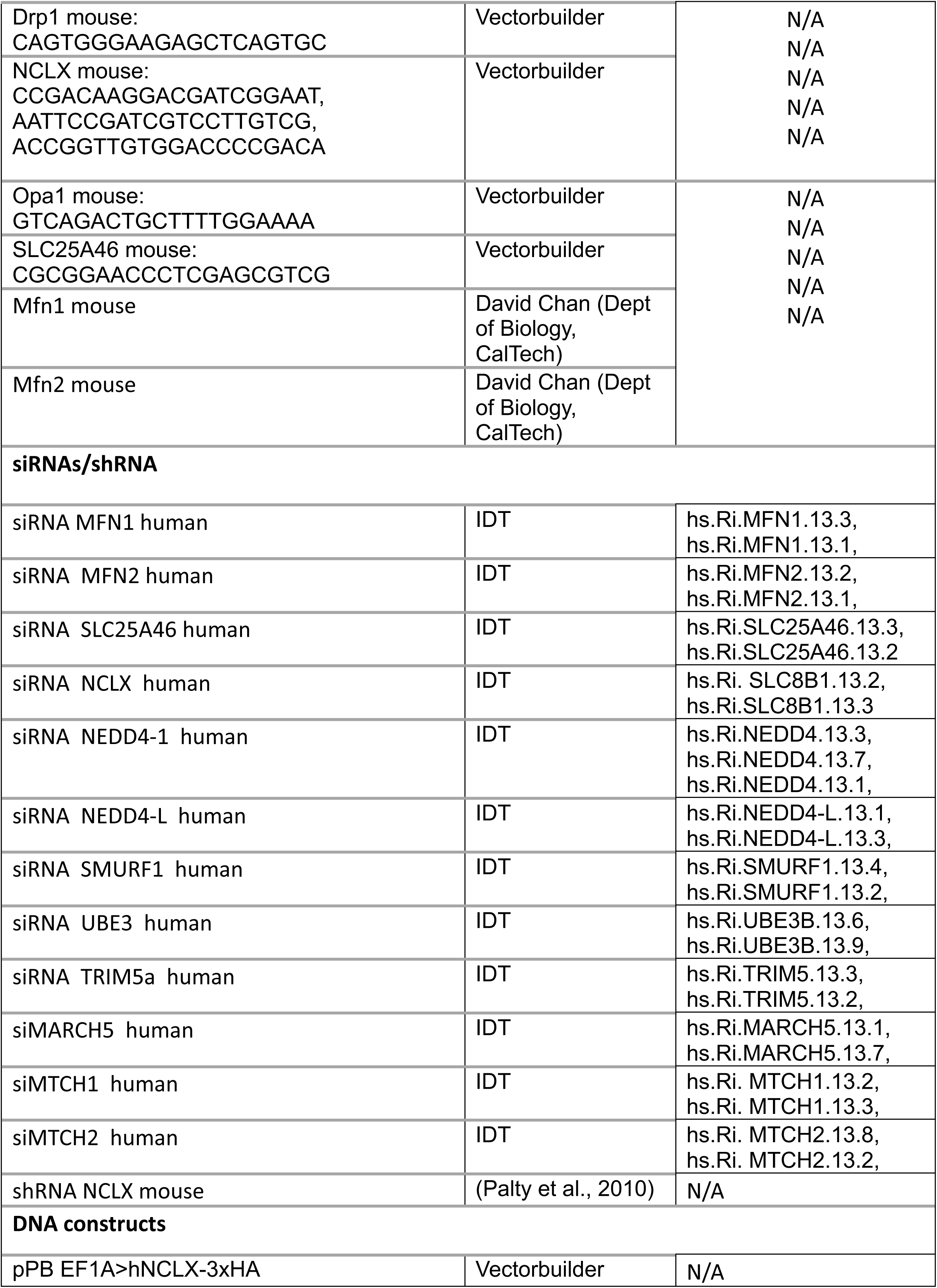
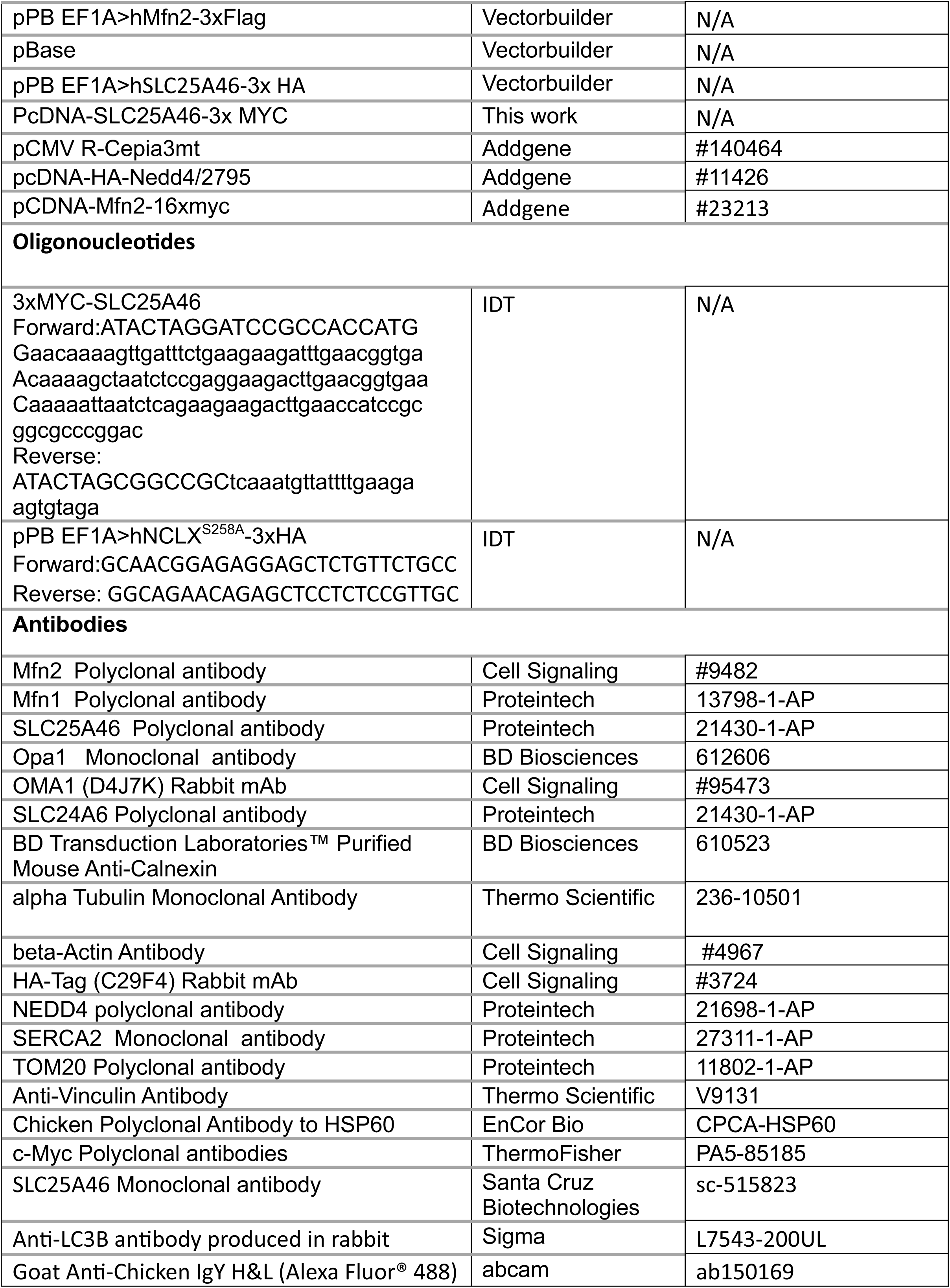
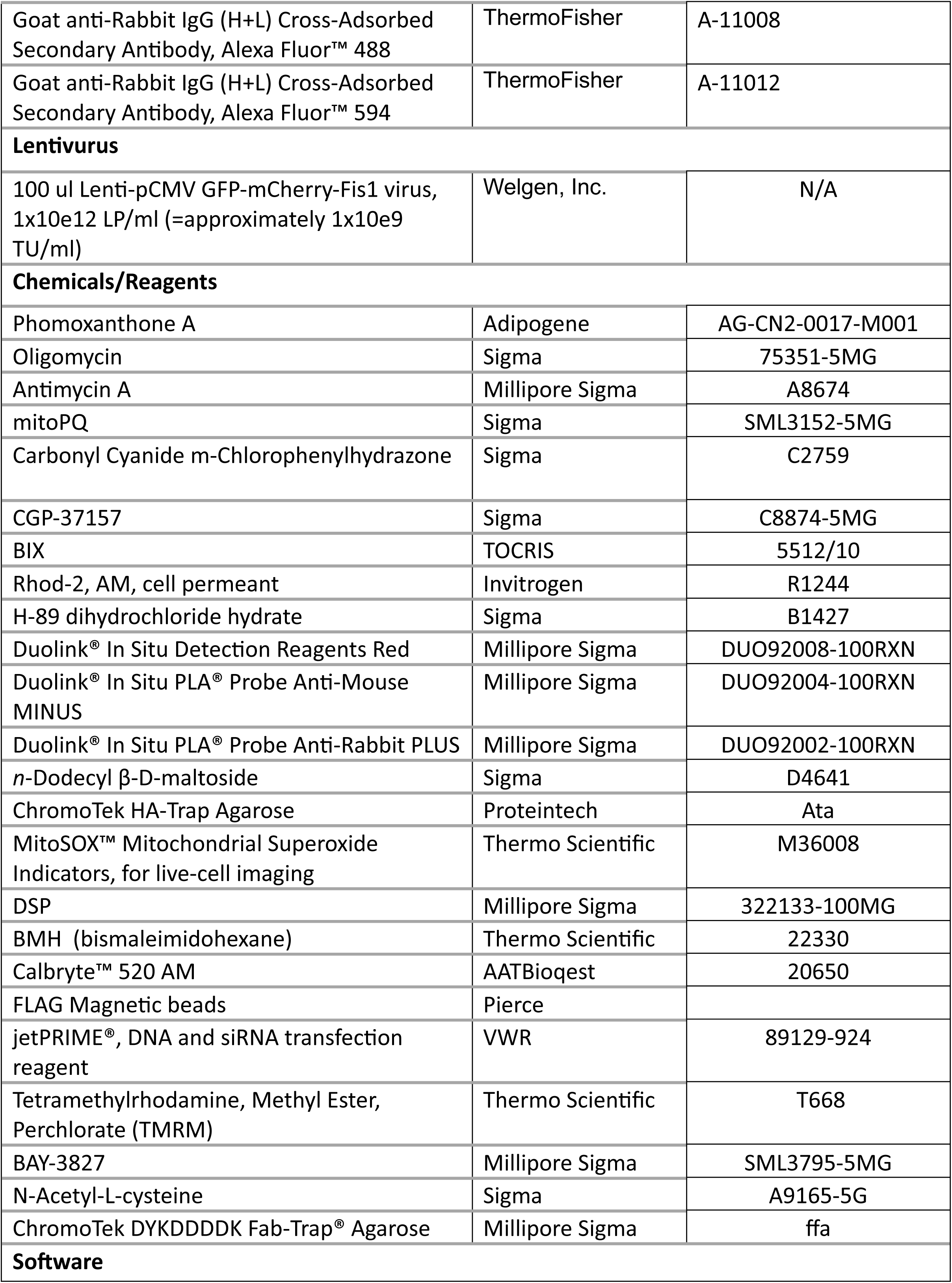
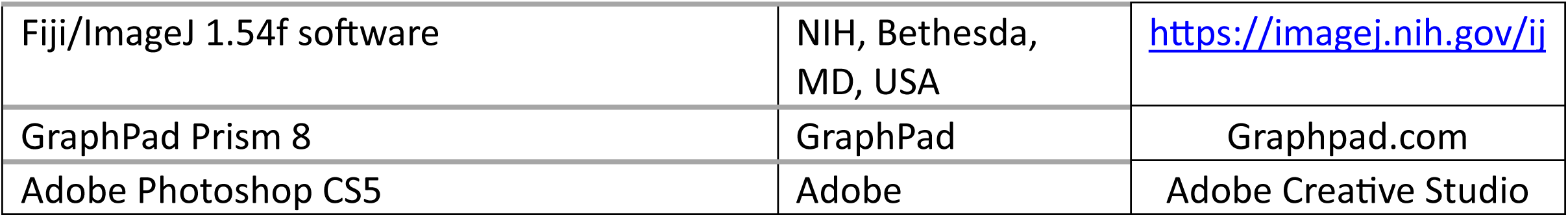

## References

Alam MS (2018) Proximity Ligation Assay (PLA). Curr Protoc Immunol 123: e58

Ali R, Parelkar SS, Thompson PR, Mitroka-Batsford S, Yerramilli S, Scarlata SF, Mistretta KS, Coburn JM, Mattson AE (2022) Phomoxanthone A Targets ATP Synthase. Chemistry 28: e202202397

Antonucci S, Mulvey JF, Burger N, Di Sante M, Hall AR, Hinchy EC, Caldwell ST, Gruszczyk AV, Deshwal S, Hartley RC et al (2019) Selective mitochondrial superoxide generation in vivo is cardioprotective through hormesis. Free Radic Biol Med 134: 678–687

Assali EA, Jones AE, Veliova M, Acin-Perez R, Taha M, Miller N, Shum M, Oliveira MF, Las G, Liesa M et al (2020) NCLX prevents cell death during adrenergic activation of the brown adipose tissue. Nat Commun 11: 3347

Baricault L, Segui B, Guegand L, Olichon A, Valette A, Larminat F, Lenaers G (2007) OPA1 cleavage depends on decreased mitochondrial ATP level and bivalent metals. Exp Cell Res 313: 3800–3808

Bohler P, Stuhldreier F, Anand R, Kondadi AK, Schlutermann D, Berleth N, Deitersen J, Wallot-Hieke N, Wu W, Frank M et al (2018) The mycotoxin phomoxanthone A disturbs the form and function of the inner mitochondrial membrane. Cell Death Dis 9: 286

Braganza A, Li J, Zeng X, Yates NA, Dey NB, Andrews J, Clark J, Zamani L, Wang XH, St Croix C et al (2017) UBE3B Is a Calmodulin-regulated, Mitochondrion-associated E3 Ubiquitin Ligase. J Biol Chem 292: 2470–2484

Braman J, Papworth C, Greener A (1996) Site-directed mutagenesis using double-stranded plasmid DNA templates. Methods Mol Biol 57: 31–44

Ceccacci S, Deitersen J, Mozzicafreddo M, Morretta E, Proksch P, Wesselborg S, Stork B, Monti MC (2020) Carbamoyl-Phosphate Synthase 1 as a Novel Target of Phomoxanthone A, a Bioactive Fungal Metabolite. Biomolecules 10

Cereghetti GM, Stangherlin A, Martins de Brito O, Chang CR, Blackstone C, Bernardi P, Scorrano L (2008) Dephosphorylation by calcineurin regulates translocation of Drp1 to mitochondria. Proceedings of the National Academy of Sciences of the United States of America 105: 15803–15808

Chakrabarti R, Ji WK, Stan RV, de Juan Sanz J, Ryan TA, Higgs HN (2018) INF2-mediated actin polymerization at the ER stimulates mitochondrial calcium uptake, inner membrane constriction, and division. J Cell Biol 217: 251–268

Chen H, Detmer SA, Ewald AJ, Griffin EE, Fraser SE, Chan DC (2003) Mitofusins Mfn1 and Mfn2 coordinately regulate mitochondrial fusion and are essential for embryonic development. J Cell Biol 160: 189–200

Chen Y, Dorn GW, 2nd (2013) PINK1-phosphorylated mitofusin 2 is a Parkin receptor for culling damaged mitochondria. Science 340: 471–475

Cribbs JT, Strack S (2007) Reversible phosphorylation of Drp1 by cyclic AMP-dependent protein kinase and calcineurin regulates mitochondrial fission and cell death. EMBO Rep 8: 939–944

Dasgupta A, Chen KH, Lima PDA, Mewburn J, Wu D, Al-Qazazi R, Jones O, Tian L, Potus F, Bonnet S et al (2021) PINK1-induced phosphorylation of mitofusin 2 at serine 442 causes its proteasomal degradation and promotes cell proliferation in lung cancer and pulmonary arterial hypertension. FASEB J 35: e21771

de Brito OM, Scorrano L (2008) Mitofusin 2 tethers endoplasmic reticulum to mitochondria. Nature 456: 605–610

De Stefani D, Patron M, Rizzuto R (2015) Structure and function of the mitochondrial calcium uniporter complex. Biochim Biophys Acta 1853: 2006–2011

De Stefani D, Rizzuto R, Pozzan T (2016) Enjoy the Trip: Calcium in Mitochondria Back and Forth. Annu Rev Biochem 85: 161–192

Dong Y, Zhuang XX, Wang YT, Tan J, Feng D, Li M, Zhong Q, Song Z, Shen HM, Fang EF et al (2023) Chemical mitophagy modulators: Drug development strategies and novel regulatory mechanisms. Pharmacol Res 194: 106835

Ehses S, Raschke I, Mancuso G, Bernacchia A, Geimer S, Tondera D, Martinou JC, Westermann B, Rugarli EI, Langer T (2009) Regulation of OPA1 processing and mitochondrial fusion by m-AAA protease isoenzymes and OMA1. J Cell Biol 187: 1023–1036

Ekhator ES, Fazzari M, Newman RH (2025) Redox Regulation of cAMP-Dependent Protein Kinase and Its Role in Health and Disease. Life (Basel) 15

Franco A, Li J, Kelly DP, Hershberger RE, Marian AJ, Lewis RM, Song M, Dang X, Schmidt AD, Mathyer ME et al (2023) A human mitofusin 2 mutation can cause mitophagic cardiomyopathy. Elife 12

Fung TS, Chakrabarti R, Higgs HN (2023) The multiple links between actin and mitochondria. Nat Rev Mol Cell Biol 24: 651–667

Garbincius JF, Elrod JW (2025) Mitochondrial sodium-calcium exchange-Can TMEM65 do it alone? Cell Metab 37: 1927–1928

Garbincius JF, Salik O, Cohen HM, Choya-Foces C, Mangold AS, Makhoul AD, Schmidt AE, Khalil DY, Doolittle JJ, Wilkinson AS et al (2025) TMEM65 regulates and is required for NCLX-dependent mitochondrial calcium efflux. Nat Metab 7: 714–729

Han F, Wu S, Dong Y, Liu Y, Sun B, Chen L (2024) Aberrant expression of NEDD4L disrupts mitochondrial homeostasis by downregulating CaMKKbeta in diabetic kidney disease. J Transl Med 22: 465

Han XJ, Lu YF, Li SA, Kaitsuka T, Sato Y, Tomizawa K, Nairn AC, Takei K, Matsui H, Matsushita M (2008) CaM kinase I alpha-induced phosphorylation of Drp1 regulates mitochondrial morphology. J Cell Biol 182: 573–585

Head B, Griparic L, Amiri M, Gandre-Babbe S, van der Bliek AM (2009) Inducible proteolytic inactivation of OPA1 mediated by the OMA1 protease in mammalian cells. J Cell Biol 187: 959–966

Hernandez-Alvarez MI, Sebastian D, Vives S, Ivanova S, Bartoccioni P, Kakimoto P, Plana N, Veiga SR, Hernandez V, Vasconcelos N et al (2019) Deficient Endoplasmic Reticulum-Mitochondrial Phosphatidylserine Transfer Causes Liver Disease. Cell 177: 881–895 e817

Hernansanz-Agustin P (2025) The unexpected role of Na(+) in mitochondrial bioenergetics, ROS production and homeostasis. Arch Biochem Biophys 772: 110544

Hernansanz-Agustin P, Morales-Vidal C, Calvo E, Natale P, Marti-Mateos Y, Jaroszewicz SN, Cabrera-Alarcon JL, Acin-Perez R, Lopez-Montero I, Vazquez J et al (2024) A transmitochondrial sodium gradient controls membrane potential in mammalian mitochondria. Cell 187: 6599–6613 e6521

Hu Y, Chen H, Zhang L, Lin X, Li X, Zhuang H, Fan H, Meng T, He Z, Huang H et al (2021) The AMPK-MFN2 axis regulates MAM dynamics and autophagy induced by energy stresses. Autophagy 17: 1142–1156

Inagaki S, Suzuki Y, Kawasaki K, Kondo R, Imaizumi Y, Yamamura H (2023) Mitofusin 1 and 2 differentially regulate mitochondrial function underlying Ca(2+) signaling and proliferation in rat aortic smooth muscle cells. Biochem Biophys Res Commun 645: 137–146

Iorio R, Celenza G, Petricca S (2021) Mitophagy: Molecular Mechanisms, New Concepts on Parkin Activation and the Emerging Role of AMPK/ULK1 Axis. Cells 11

Islam MM, Takeuchi A, Matsuoka S (2020) Membrane current evoked by mitochondrial Na(+)-Ca(2+) exchange in mouse heart. J Physiol Sci 70: 24

Karbowski M, Lee YJ, Gaume B, Jeong SY, Frank S, Nechushtan A, Santel A, Fuller M, Smith CL, Youle RJ (2002) Spatial and temporal association of Bax with mitochondrial fission sites, Drp1, and Mfn2 during apoptosis. J Cell Biol 159: 931–938

Kleele T, Rey T, Winter J, Zaganelli S, Mahecic D, Perreten Lambert H, Ruberto FP, Nemir M, Wai T, Pedrazzini T et al (2021) Distinct fission signatures predict mitochondrial degradation or biogenesis. Nature 593: 435–439

Kolitsida P, Zhou J, Rackiewicz M, Nolic V, Dengjel J, Abeliovich H (2019) Phosphorylation of mitochondrial matrix proteins regulates their selective mitophagic degradation. Proc Natl Acad Sci U S A 116: 20517–20527

Korobova F, Gauvin TJ, Higgs HN (2014) A role for myosin II in mammalian mitochondrial fission. Curr Biol 24: 409–414

Korobova F, Ramabhadran V, Higgs HN (2013) An actin-dependent step in mitochondrial fission mediated by the ER-associated formin INF2. Science 339: 464–467

Kostic M, Katoshevski T, Sekler I (2018) Allosteric Regulation of NCLX by Mitochondrial Membrane Potential Links the Metabolic State and Ca(2+) Signaling in Mitochondria. Cell Rep 25: 3465–3475 e3464

Kostic M, Ludtmann MH, Bading H, Hershfinkel M, Steer E, Chu CT, Abramov AY, Sekler I (2015) PKA Phosphorylation of NCLX Reverses Mitochondrial Calcium Overload and Depolarization, Promoting Survival of PINK1-Deficient Dopaminergic Neurons. Cell Rep 13: 376–386

Kostic M, Sekler I (2019) Functional properties and mode of regulation of the mitochondrial Na(+)/Ca(2+) exchanger, NCLX. Semin Cell Dev Biol 94: 59–65

Kumar AV, Mills J, Lapierre LR (2022) Selective Autophagy Receptor p62/SQSTM1, a Pivotal Player in Stress and Aging. Front Cell Dev Biol 10: 793328

Labbe K, Murley A, Nunnari J (2014) Determinants and functions of mitochondrial behavior. Annu Rev Cell Dev Biol 30: 357–391

Lin Q, Dai Q, Meng H, Sun A, Wei J, Peng K, Childress C, Chen M, Shao G, Yang W (2017) The HECT E3 ubiquitin ligase NEDD4 interacts with and ubiquitylates SQSTM1 for inclusion body autophagy. J Cell Sci 130: 3839–3850

Lu K, Li P, Zhang M, Xing G, Li X, Zhou W, Bartlam M, Zhang L, Rao Z, He F (2011) Pivotal role of the C2 domain of the Smurf1 ubiquitin ligase in substrate selection. J Biol Chem 286: 16861–16870

MacVicar TD, Lane JD (2014) Impaired OMA1-dependent cleavage of OPA1 and reduced DRP1 fission activity combine to prevent mitophagy in cells that are dependent on oxidative phosphorylation. J Cell Sci 127: 2313–2325

Martinez M, Martinez NA, Silva WI (2017) Measurement of the Intracellular Calcium Concentration with Fura-2 AM Using a Fluorescence Plate Reader. Bio Protoc 7: e2411

Montero M, Alonso MT, Albillos A, Garcia-Sancho J, Alvarez J (2001) Mitochondrial Ca(2+)-induced Ca(2+) release mediated by the Ca(2+) uniporter. Mol Biol Cell 12: 63–71

Mourier A, Motori E, Brandt T, Lagouge M, Atanassov I, Galinier A, Rappl G, Brodesser S, Hultenby K, Dieterich C et al (2015) Mitofusin 2 is required to maintain mitochondrial coenzyme Q levels. J Cell Biol 208: 429–442

Narendra D, Tanaka A, Suen DF, Youle RJ (2008) Parkin is recruited selectively to impaired mitochondria and promotes their autophagy. J Cell Biol 183: 795–803

Numata M, Petrecca K, Lake N, Orlowski J (1998) Identification of a mitochondrial Na+/H+ exchanger. J Biol Chem 273: 6951–6959

Palty R, Silverman WF, Hershfinkel M, Caporale T, Sensi SL, Parnis J, Nolte C, Fishman D, Shoshan-Barmatz V, Herrmann S et al (2010) NCLX is an essential component of mitochondrial Na+/Ca2+ exchange. Proc Natl Acad Sci U S A 107: 436–441

Peng H, Yang J, Li G, You Q, Han W, Li T, Gao D, Xie X, Lee BH, Du J et al (2017) Ubiquitylation of p62/sequestosome1 activates its autophagy receptor function and controls selective autophagy upon ubiquitin stress. Cell Res 27: 657–674

Peng W, Wong YC, Krainc D (2020) Mitochondria-lysosome contacts regulate mitochondrial Ca(2+) dynamics via lysosomal TRPML1. Proc Natl Acad Sci U S A 117: 19266–19275

Plant PJ, Lafont F, Lecat S, Verkade P, Simons K, Rotin D (2000) Apical membrane targeting of Nedd4 is mediated by an association of its C2 domain with annexin XIIIb. J Cell Biol 149: 1473–1484

Quiros PM, Ramsay AJ, Sala D, Fernandez-Vizarra E, Rodriguez F, Peinado JR, Fernandez-Garcia MS, Vega JA, Enriquez JA, Zorzano A et al (2012) Loss of mitochondrial protease OMA1 alters processing of the GTPase OPA1 and causes obesity and defective thermogenesis in mice. The EMBO journal 31: 2117–2133

Rainbolt TK, Lebeau J, Puchades C, Wiseman RL (2016) Reciprocal Degradation of YME1L and OMA1 Adapts Mitochondrial Proteolytic Activity during Stress. Cell Rep 14: 2041–2049

Saha B, Olsvik H, Williams GL, Oh S, Evjen G, Sjottem E, Mandell MA (2024) TBK1 is ubiquitinated by TRIM5alpha to assemble mitophagy machinery. Cell Rep 43: 114294

Samanta K, Mirams GR, Parekh AB (2018) Sequential forward and reverse transport of the Na(+) Ca(2+) exchanger generates Ca(2+) oscillations within mitochondria. Nat Commun 9: 156

Santel A, Fuller MT (2001) Control of mitochondrial morphology by a human mitofusin. J Cell Sci 114: 867–874

Schuettpelz J, Janer A, Antonicka H, Shoubridge EA (2023) The role of the mitochondrial outer membrane protein SLC25A46 in mitochondrial fission and fusion. Life Sci Alliance 6

Sebastian D, Hernandez-Alvarez MI, Segales J, Sorianello E, Munoz JP, Sala D, Waget A, Liesa M, Paz JC, Gopalacharyulu P et al (2012) Mitofusin 2 (Mfn2) links mitochondrial and endoplasmic reticulum function with insulin signaling and is essential for normal glucose homeostasis. Proc Natl Acad Sci U S A 109: 5523–5528

Steffen J, Vashisht AA, Wan J, Jen JC, Claypool SM, Wohlschlegel JA, Koehler CM (2017) Rapid degradation of mutant SLC25A46 by the ubiquitin-proteasome system results in MFN1/2-mediated hyperfusion of mitochondria. Mol Biol Cell 28: 600–612

Stepp WL, Tortarolo G, Landoni JC, Durmus EB, Rodriguez Alvarez SN, Douglass KM, Weigert M, Manley S (2026) Smart hybrid microscopy for cell-friendly detection of rare events. Nat Commun

Sun A, Wei J, Childress C, Shaw JHt, Peng K, Shao G, Yang W, Lin Q (2017) The E3 ubiquitin ligase NEDD4 is an LC3-interactive protein and regulates autophagy. Autophagy 13: 522–537

Taha M, Assali EA, Ben-Kasus Nissim T, Stutzmann GE, Shirihai OS, Hershfinkel M, Sekler I (2024) NCLX controls hepatic mitochondrial Ca(2+) extrusion and couples hormone-mediated mitochondrial Ca(2+) oscillations with gluconeogenesis. Mol Metab 87: 101982

Tondera D, Grandemange S, Jourdain A, Karbowski M, Mattenberger Y, Herzig S, Da Cruz S, Clerc P, Raschke I, Merkwirth C et al (2009) SLP-2 is required for stress-induced mitochondrial hyperfusion. EMBO J 28: 1589–1600

Toyama EQ, Herzig S, Courchet J, Lewis TL, Jr., Loson OC, Hellberg K, Young NP, Chen H, Polleux F, Chan DC et al (2016) Metabolism. AMP-activated protein kinase mediates mitochondrial fission in response to energy stress. Science 351: 275–281

Twig G, Elorza A, Molina AJ, Mohamed H, Wikstrom JD, Walzer G, Stiles L, Haigh SE, Katz S, Las G et al (2008) Fission and selective fusion govern mitochondrial segregation and elimination by autophagy. EMBO J 27: 433–446

Verhoeven K, Claeys KG, Zuchner S, Schroder JM, Weis J, Ceuterick C, Jordanova A, Nelis E, De Vriendt E, Van Hul M et al (2006) MFN2 mutation distribution and genotype/phenotype correlation in Charcot-Marie-Tooth type 2. Brain : a journal of neurology 129: 2093–2102

Vetralla M, Wischhof L, Kahsay A, Cadenelli V, Scifo E, Xie B, Sbrissa M, Habert MS, Ehninger D, Rizzuto R et al (2025) TMEM65-dependent Ca2+ extrusion safeguards mitochondrial homeostasis. Nat Commun

Wang N, Wang X, Lan B, Gao Y, Cai Y (2025a) DRP1, fission and apoptosis. Cell Death Discov 11: 150

Wang Q, Sun Y, Li TY, Auwerx J (2026) Mitophagy in the pathogenesis and management of disease. Cell Res 36: 11–37

Wang R, Wang J, Yu J, Li Z, Zhang M, Chen Y, Liu F, Jiang D, Guo J, Li X et al (2025b) Mfn2 regulates calcium homeostasis and suppresses PASMCs proliferation via interaction with IP3R3 to mitigate pulmonary arterial hypertension. J Transl Med 23: 366

Wilkins HM, Troutwine BR, Menta BW, Manley SJ, Strope TA, Lysaker CR, Swerdlow RH (2022) Mitochondrial Membrane Potential Influences Amyloid-beta Protein Precursor Localization and Amyloid-beta Secretion. J Alzheimers Dis 85: 381–394

Wong YC, Kim S, Peng W, Krainc D (2019) Regulation and Function of Mitochondria-Lysosome Membrane Contact Sites in Cellular Homeostasis. Trends Cell Biol 29: 500–513

Wong YC, Ysselstein D, Krainc D (2018) Mitochondria-lysosome contacts regulate mitochondrial fission via RAB7 GTP hydrolysis. Nature 554: 382–386

Xiao S, Yu Y, Liao M, Song D, Xu X, Tian L, Zhang R, Lyu H, Guo D, Zhang Q et al (2025) Post-Translational Modification of p62: Roles and Regulations in Autophagy. Cells 14

Xu S, Wang P, Zhang H, Gong G, Gutierrez Cortes N, Zhu W, Yoon Y, Tian R, Wang W (2016) CaMKII induces permeability transition through Drp1 phosphorylation during chronic beta-AR stimulation. Nat Commun 7: 13189

Yang JF, Xing X, Luo L, Zhou XW, Feng JX, Huang KB, Liu H, Jin S, Liu YN, Zhang SH et al (2023) Mitochondria-ER contact mediated by MFN2-SERCA2 interaction supports CD8(+) T cell metabolic fitness and function in tumors. Sci Immunol 8: eabq2424

Youle RJ, van der Bliek AM (2012) Mitochondrial fission, fusion, and stress. Science 337: 1062–1065

Zaninello M, Palikaras K, Sotiriou A, Tavernarakis N, Scorrano L (2022) Sustained intracellular calcium rise mediates neuronal mitophagy in models of autosomal dominant optic atrophy. Cell Death Differ 29: 167–177

Zhang J, Wang X, Vikash V, Ye Q, Wu D, Liu Y, Dong W (2016) ROS and ROS-Mediated Cellular Signaling. Oxid Med Cell Longev 2016: 4350965

Zhang JL, Chang YC, Lai PH, Yeh HI, Tsai CW, Huang YL, Liu TY, Lee IC, Foulon N, Xu Y et al (2025) TMEM65 functions as the mitochondrial Na(+)/Ca(2+) exchanger. Nat Cell Biol 27: 1301–1310

Zhou W, Chen KH, Cao W, Zeng J, Liao H, Zhao L, Guo X (2010) Mutation of the protein kinase A phosphorylation site influences the anti-proliferative activity of mitofusin 2. Atherosclerosis 211: 216–223

Zuchner S, Mersiyanova IV, Muglia M, Bissar-Tadmouri N, Rochelle J, Dadali EL, Zappia M, Nelis E, Patitucci A, Senderek J et al (2004) Mutations in the mitochondrial GTPase mitofusin 2 cause Charcot-Marie-Tooth neuropathy type 2A. Nat Genet 36: 449–451

